# Unlocking Targeted Underwater Delivery of Coral Probiotics

**DOI:** 10.64898/2026.11.04.764210

**Authors:** Martina Pierina Facchinelli, Marco Contardi, Jessica Caldwell, Fabrizio Fiorentini, Isabella Ghizzi, Vittoria Calabretta, Luca Ceseracciu, Tania Pomili, Valerio Isa, Gabriele Corigliano, Camilla Rinaldi, Vincenzo Scribano, Elena Bellini, Lorenzo Ravelli, Silvia Lavorano, Isabella Gandolfi, Paolo Galli, Simone Montano, Athanassia Athanassiou

**Affiliations:** Smart Materials, Istituto Italiano di Tecnologia, Via Morego 30, 16163, Genova, Italy; Department of Earth and Environmental Sciences (DISAT), University of Milano-Bicocca, Piazza della Scienza 1, 20126, Milan, Italy; Translational Pharmacology Facility, Istituto Italiano di Tecnologia, Via Morego 30, 16163, Genova, Italy; Materials Characterization Facility, Istituto Italiano di Tecnologia, Via Morego 30, 16163, Genova, Italy; Electron Microscopy Facility, Istituto Italiano di Tecnologia, Via Morego 30, 16163, Genova, Italy; Center of Nanoscience and Technology, Via Raffaele Rubattino 81, 20134, Milan, Italy; Costa Edutainment SpA - Acquario di Genova, 16128, Genova, Italy; MaRHE Center (Marine Research and High Education Center), Magoodhoo Island, Faafu Atoll 12030, Republic of Maldives

**Keywords:** microbial therapy, underwater delivery systems, beads, hydrogels, filaments, *Pseudoalteromonas*

## Abstract

The rapid global decline of coral reefs has intensified the need for innovative conservation strategies, including microbial therapies to enhance coral health and resilience. A major limitation, however, is the lack of effective delivery systems for administering Beneficial Microorganisms for Corals (BMCs). In this study, we present natural, biodegradable systems for the underwater delivery of probiotic bacteria to corals. Three customized alginate-based delivery formats (beads, hydrogels, and filaments) were developed and loaded with the probiotic *Pseudoalteromonas peptidolytica*. The materials were loaded with the bacterium at an average concentration of 10^4^ CFU/g of dry polymer and engineered to sink underwater while keeping their structural integrity over time. *In vitro* assays showed controlled release kinetics, with encapsulated probiotics remaining viable and capable of replication. *In vivo* experiments compared syringe-based inoculation with the application of alginate beads, hydrogels, and filaments to corals. Microbiome analyses performed before and after inoculation confirmed successful probiotic colonization following deployment of the alginate-based delivery systems. Relative abundance of the probiotic genus *Pseudoalteromonas* increased markedly in treated corals, reaching 11.5 % after 48 h with alginate beads, 12.7 % after 48 h with hydrogels, and 12.6 % after 24 h with filaments. Furthermore, the degradation mechanisms of the three delivery systems were investigated. As the first biomaterial-based underwater delivery systems specifically engineered for coral probiotics, these alginate-based materials provide a promising foundation for advancing microbial therapies in both controlled aquaria and natural reef environments.

## 1. Introduction

One of the most known and studied symbioses in the marine environment is the one between corals and photosynthetic dinoflagellates of the family Symbiodiniaceae, also known as zooxanthellae, which provide the energy required by the coral for metabolic activity (*1–4*). However, corals can engage in many other symbioses with microorganisms, including bacteria, fungi, viruses, and archaea, forming the so-called holobiont (*5, 6*). Specifically, the total community of microorganisms associated with the coral is defined as the microbiome (*7*), and some of these microorganisms are species-specific or shared across multiple coral taxa, thereby constituting a microbial fingerprint that uniquely characterizes each coral colony (*6, 8*). The recognition of bacterial functions and roles in coral health led to the development of the Coral Probiotic Hypothesis (CPH). This hypothesis suggests that, under changing environmental conditions, the microbial composition of the coral holobiont can be modified, enabling the coral to adapt to new conditions (*9*). Within the coral microbiome, the microbial community is highly heterogeneous (*10–12*), and each bacterial member may play a crucial role in maintaining coral health and homeostasis. While the functions of some microorganisms have been characterized, the ecological roles of many others remain unknown (*13*). In addition, certain opportunistic bacteria can exhibit context-dependent behavior, shifting between beneficial and potentially detrimental interactions according to environmental conditions and the level of stress experienced by the coral host (*5, 13–15*). BMCs refer to all probiotic microorganisms that can benefit coral health (*16*). They can contribute substantially to energy metabolism and nutrient acquisition, larval settlement and recruitment, and detoxification processes, helping to mitigate or transform potentially harmful compounds within the holobiont microenvironment (*16–30*). Furthermore, some bacteria, such as *Pseudoalteromonas* spp. and *Roseobacter* spp., can play an important role in host defense by limiting the proliferation of opportunistic or pathogenic microorganisms through the production of antimicrobial compounds and competitive interactions within the microbiome (*31–37*).

Despite these reported beneficial effects, several key aspects remain insufficiently understood. Critical knowledge gaps include optimal delivery routes, probiotic persistence within the coral microbiome, the number of inoculations required to induce significant shifts in the microbiome, and the resulting dynamics of introduced strains within the native microbial community. Regarding the administration and delivery routes of BMCs, most studies performed in aquaria and in the field rely on repeated applications using a syringe loaded with probiotic consortia, typically two or more inoculations, reaching a maximum of three per week for three months (*38–47*), making these approaches highly time-consuming and potentially impractical for large-scale reef interventions. Moreover, even with repeated inoculations, the final relative abundance of the introduced probiotics on corals often remains low, generally ranging from ∼0.1% to ∼4% of the total microbial community (*41, 42, 46*). In some cases, higher levels have been reported, with relative abundances reaching ∼8% (*39, 40*) and up to 10–20% after seven inoculations (*44*), while concentrations near 60% have been observed in one study after only four applications (*47*). Even though studies using a single probiotic inoculation in closed systems such as aquaria or tanks also exist, the relative abundance of the delivered probiotic detected on the coral tissue is less than 5% (*48–50*). Similarly, BMC inoculation has been performed in the sea using a syringe containing a bacterial consortium, resulting in probiotic establishment in the microbiome of up to 2.5% (*42*). Nevertheless, several complicating factors must be considered when delivering microorganisms by injecting bacterial solutions through a syringe. These include their direct dilution in water, interactions with the diverse microorganisms inhabiting the marine environment, and the influence of waves and currents, which can cause uncontrolled diffusion and hinder delivery (*51*).

Another proposed delivery strategy is the use of living carriers, such as rotifers, tested as vectors for successfully delivering probiotics to corals. However, the concentration of probiotics effectively delivered to the corals was not quantified (*52*). These observations highlight the need to develop strategies that minimize the number of inoculations while enabling an effective delivery of the probiotic to the coral microbiome.

An ideal strategy could be to design an underwater delivery system to properly administrate probiotics to corals. The vehicle should remain attached to coral structures, allowing targeted release of BMCs without immediately dispersing the consortium into the surrounding environment or triggering a coral stress response due to material-related toxicity, and be eco-friendly. Biocompatible, biodegradable, and non-toxic polymers have been selected to produce probiotic delivery systems in other fields, such as biomedicine (*53, 54*). For developing an underwater probiotic delivery system (*51*), chitin, agar, and alginate have been proposed as viable options, however, to our knowledge, their effectiveness in successfully delivering BMCs to corals, and the subsequent effects on the coral-associated microbiome remain an open question and challenge to be addressed.

In this work, the anionic polymer alginate was specifically selected as the delivery material because it is naturally present in brown algae (*55*), can be processed under mild conditions, has a simple structure, low toxicity and exhibits excellent physicochemical properties, while it can easily form a gel matrix for the bacteria (*56–58*). In addition, alginate is a major component of the bacterial biofilm matrix, forming a scaffold that provides porosity, mechanical stability, and water retention (*59*). Its structure enables microorganisms to communicate and interact, thereby forming synergistic microconsortia (*60*). The alginate matrix supplies nutrients to biofilm-associated organisms and protects them from desiccation, antibiotics, and ultraviolet radiation (*59, 60*). Additionally, alginate has already been proposed as a polymer for probiotic encapsulation in both human biomedical treatments and marine applications (*16, 43, 51, 58, 61–64*).

Among the BMC strains, *Pseudoalteromonas* species, a genus of Gram-negative bacteria, have been shown to benefit coral health when used as part of probiotic consortia inoculated into corals (*38–40, 42–44, 47, 65*). Their physiological abundance has been reported to be usually less than 1% (*66*). Previous investigations have shown that even following treatment, *Pseudoalteromonas* sp. does not persist in coral tissues at high abundance, typically accounting for only ∼0.5–2.5% of the total microbiome.

For this work, three customized alginate delivery systems (beads, hydrogels, and filaments) were developed and loaded with the probiotic bacterium *Pseudoalteromonas peptidolytica*. The BMC loading was quantified, and the chemical and mechanical properties of the delivery systems were analyzed. To assess the impact of encapsulation on probiotic physiology, the kinetics of probiotic release were studied *in vitro*. The three delivery platforms were then tested on three coral species, with different 3D morphologies, to assess the strategy’s versatility. Under controlled aquarium conditions, probiotic delivery and changes in coral-associated microbiome composition were monitored at different time points over 72 h. The performance of our three alginate-based delivery systems was compared with that of conventional syringe-based probiotic administration, the most widely reported delivery method in the literature. Finally, physicochemical changes occurring during biodegradation and the microbial communities associated with this process were investigated using energy-dispersive X-ray spectroscopy (EDX) and microbiome profiling, respectively.

## 2. Results

### 2.1 Fabrication, encapsulation efficiency, and morphology of the bacteria-loaded materials

A schematic representation of the composition of the materials, the potential interaction between the materials and the corals, and the application in situ is described in **Figure 1A**. Photographs of the beads, hydrogels, and filaments fabricated with the same alginate concentration and loaded with the probiotic *Pseudoalteromonas peptidolytica* are shown in **Figures 1B, C**, and **D**. An alginate concentration of 5% (w/v) in Marine Broth (MB) crosslinked with CaCl₂ was chosen for all systems to ensure a stable gel-like structure and a density higher than seawater (typically: 1.023 g/cm^3^ at 25°C and 1.022 g/cm^3^ at 28°C (*67, 68*)) thereby causing the materials to sink. For further details on material fabrication, see Section 5.2.

**Figure 1.**
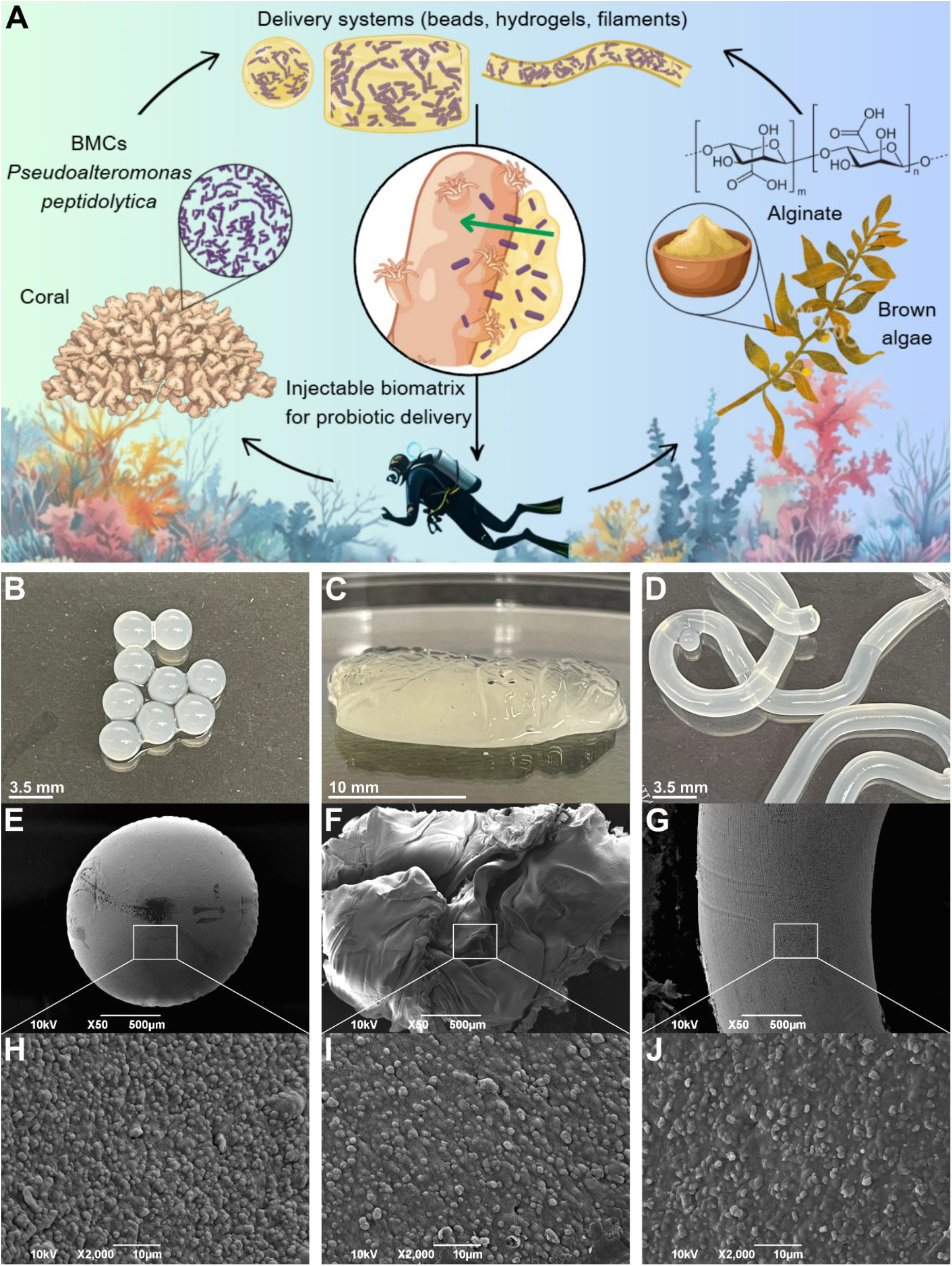
Composition and morphological characterization. (A) Schematic illustration of the composition of the three alginate-based delivery systems, the potential interaction among the materials, corals, and the environment, and the potential administration strategy. (B, C, and D) Photographs of the final loaded materials. (E, F, and G) Top-view SEM images at low magnification of loaded beads, hydrogels, and filaments, respectively. (H, I, and J) Top-view SEM images, at increased magnification, showing the surface morphologies and presence of encapsulated probiotics in loaded beads, hydrogels, and filaments, respectively.

The encapsulation efficiency of the three systems was quantified by normalizing the number of CFUs released upon disintegration of the freshly fabricated samples (*62*) to their respective dry weight, thereby expressing the results as CFUs/g of dry polymer (∼10^4^ CFU/g dry polymer). More details on the calculations are provided in **Figure S1 and Table S1**. Moreover, encapsulation efficiency was unaffected by variations in crosslinking time or CaCl₂ concentration, indicating that the encapsulation conditions were non-toxic to the probiotics.

Representative Scanning Electron Microscopy (SEM) top-view images of the loaded beads, hydrogels, and filaments are shown in **Figures 1E, F**, and **G**, respectively. At the macroscale, the loaded beads and filaments exhibited smooth, flat surfaces, whereas the hydrogels showed a higher surface roughness, likely due to the manufacturing process. At higher magnification (**Figures 1H, I**, and **J**), bacteria were visible on all surfaces. Further investigation of the cross-section in beads (**Figure S2A**) clearly revealed bacterial cells embedded within the material’s internal structure. This morphological evidence supports the effective entrapment of the probiotics within the matrix. Control SEM images showing the pristine probiotic *Pseudoalteromonas peptidolytica* are shown in **Figures S2B** and **C**, while top-view SEM images at high magnifications of blank beads, hydrogels, and filaments, all exhibiting smooth surfaces due to the absence of probiotics, are shown in **Figures S2D**, **E**, and **F**.

### 2.3 Mechanical properties

Mechanical analyses were performed to investigate the influence of crosslinking time on the overall mechanical behavior and resistance of beads, hydrogels, and filaments with and without probiotics after immersion in seawater. To assess this, nanoindentation, compression, and rheological analyses were conducted (**Figure S3A**, **S3B**, and **S3C**).

Results from nanoindentation on beads, hydrogels, and filaments are reported in **Figures 2A, B, C**, and **Table S2**, and were extrapolated from load-displacement curves (**Figures S3D** and **S3E**). Among the blank formulations, beads, hydrogels, and filaments crosslinked for 20 min exhibited the highest Young’s modulus (E) of 74.7 ± 6.4 kPa, 56.1 ± 7.7 kPa, and 53.5 ± 7.5 kPa, respectively. The E values were lower for all the probiotic-loaded systems crosslinked at 20 min, with maximum values of 61.8 ± 6.3 kPa for the beads, 39.7 ± 14.7 kPa for the hydrogels, and 49.2 ± 2.6 kPa for the filaments. No significant differences in E were detected among the crosslinking times for either the blank or the loaded systems, with all values remaining within the same order of magnitude (in kPa). Considering both these findings and the preliminary floating properties observed, a crosslinking time of 20 min was selected for subsequent experiments.

**Figure 2.**
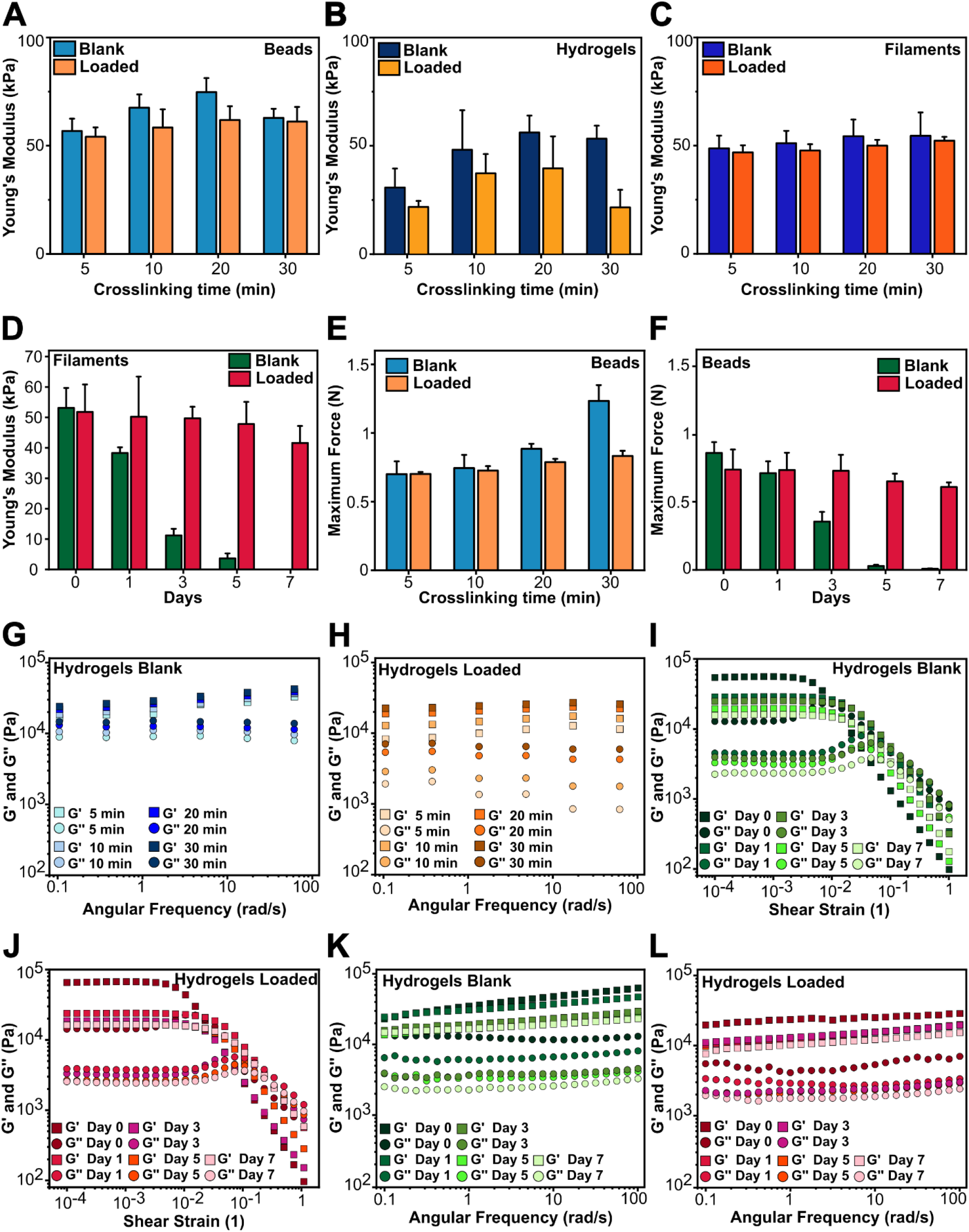
Mechanical characterization of the materials. (A, B, C) Results of nanoindentation for blank and loaded beads, hydrogels, and filaments, respectively. (D) Nanoindentation results for blank and loaded filaments developed after a crosslinking time of 20 min and immersed in seawater for 7 days. (E, F) Results of the compression test for blank and loaded beads, expressed as a function of the maximum force at break, evaluating the different crosslinking times and immersion for 7 days, respectively. (G, J) Frequency sweep test performed on blank and loaded hydrogels at different crosslinking times. (H, I) Results of the amplitude sweep test on blank and loaded hydrogels immersed in seawater for 7 days. (K, L) Frequency sweep test results on blank and loaded hydrogels immersed in seawater for 7 days.

Next, the E of the filaments was measured daily over a 7-day seawater immersion period to evaluate their stability in this medium (**Figure 2D** and **Table S3**). For the blank filaments, the E values decreased markedly up to the 6^th^ day, after which further measurements could not be obtained due to structural degradation. In contrast, the probiotic-loaded filaments maintained relatively stable E values throughout the 7-day period, showing a small reduction of approximately 19.7%.

The compression test results for the crosslinking times in beads are reported in **Figure 2E** and **Table S4**. Blank beads after 30-minute crosslinking time showed the highest maximum force at break (MFB) value of ≈ 1.23 ± 0.11 N, while the MFB values were ≈ 0.88 ± 0.03 N, ≈ 0.74 ± 0.09 N, and ≈ 0.69 ± 0.09 N for blank beads crosslinked at 20, 10, and 5 min. These results are comparable to previous compressive test studies performed on alginate beads (*69*). On the other hand, loaded beads exhibited lower stiffness and strength than blank beads, with MFB values of ≈ 0.83 ± 0.03 N, ≈ 0.78 ± 0.02 N, ≈ 0.72 ± 0.03 N, and ≈ 0.70 ± 0.01 N for beads crosslinked for 30, 20, 10, and 5 min, respectively.

In the long-term immersion experiment, as shown in **Figure 2F** and **Table S5**, the MFB of the blank beads crosslinked for 20 min decreased over time, indicating progressive structural weakening. In contrast, probiotic-loaded beads crosslinked for 20 min and immersed in seawater exhibited stable MFB values, with only a 17% reduction observed throughout the same period. This trend of increased mechanical stability due to the presence of probiotics within the alginate matrix is consistent with that observed for the filaments.

Hydrogels were characterized by rheology. Analysis of the amplitude sweep tests (**Figures S4A**, **S4B**, and **Table S6**) revealed that within the Linear Viscoelastic Region, both blank hydrogels and hydrogels loaded with probiotics exhibited solid-gel-like behavior, as indicated by a higher Storage Modulus (G’) compared to the Loss Modulus (G’’). The frequency-sweep tests further corroborated these observations: G’ was higher than G’’ across the evaluated frequency range for all the investigated crosslinking times (**Figures 2G, H**, and **Table S7**). Throughout the 7-day seawater immersion analysis, the hydrogels consistently exhibited solid-gel behavior, as evidenced by both amplitude-sweep tests within the Linear Viscoelastic Region (**Figures 2I, J**, and **Table S8**) and frequency-sweep measurements (**Figures 2K, L**, and **Table S9**). During the amplitude sweep test, after the first day, both the storage modulus (G’) and the loss modulus (G’’) decreased noticeably, indicating a structural relaxation of the material. This initial drop reflected the rearrangement or partial breakdown of the internal network formed right after sample crosslinking (*55*). After this equilibration phase, the values of G’ and G’’ remained nearly constant over the following days, suggesting that the material had reached a stable structure and no further significant changes occurred. Rheological measurements confirmed that the hydrogels exhibited quite stable behavior across all tested samples, and the results were consistent with values reported in the literature (*70*). Notably, the blank samples remained intact until day 7, indicating that the hydrogel matrix exhibits greater stability in seawater than the other two delivery systems. This result may also be attributable to their thicker shape, which likely contributed to their longer persistence.

### 2.4 Probiotic release *in vitro*

Probiotic release was assessed *in vitro* to investigate the mechanisms and kinetics of bacterial diffusion and growth following immersion in an aqueous medium; the experimental design is shown in **Figure S5**. Specifically, loaded alginate systems were placed in well plates and immersed in culture medium. At each time point (0, 1, 2, 4, 6, 8, 24, and 48 hours), an aliquot of the medium was collected and plated to quantify the bacteria released. The results for the 48-hour probiotic release from beads, hydrogels, and filaments are shown in **Figures 3A, B**, and **C**, respectively.

**Figure 3.**
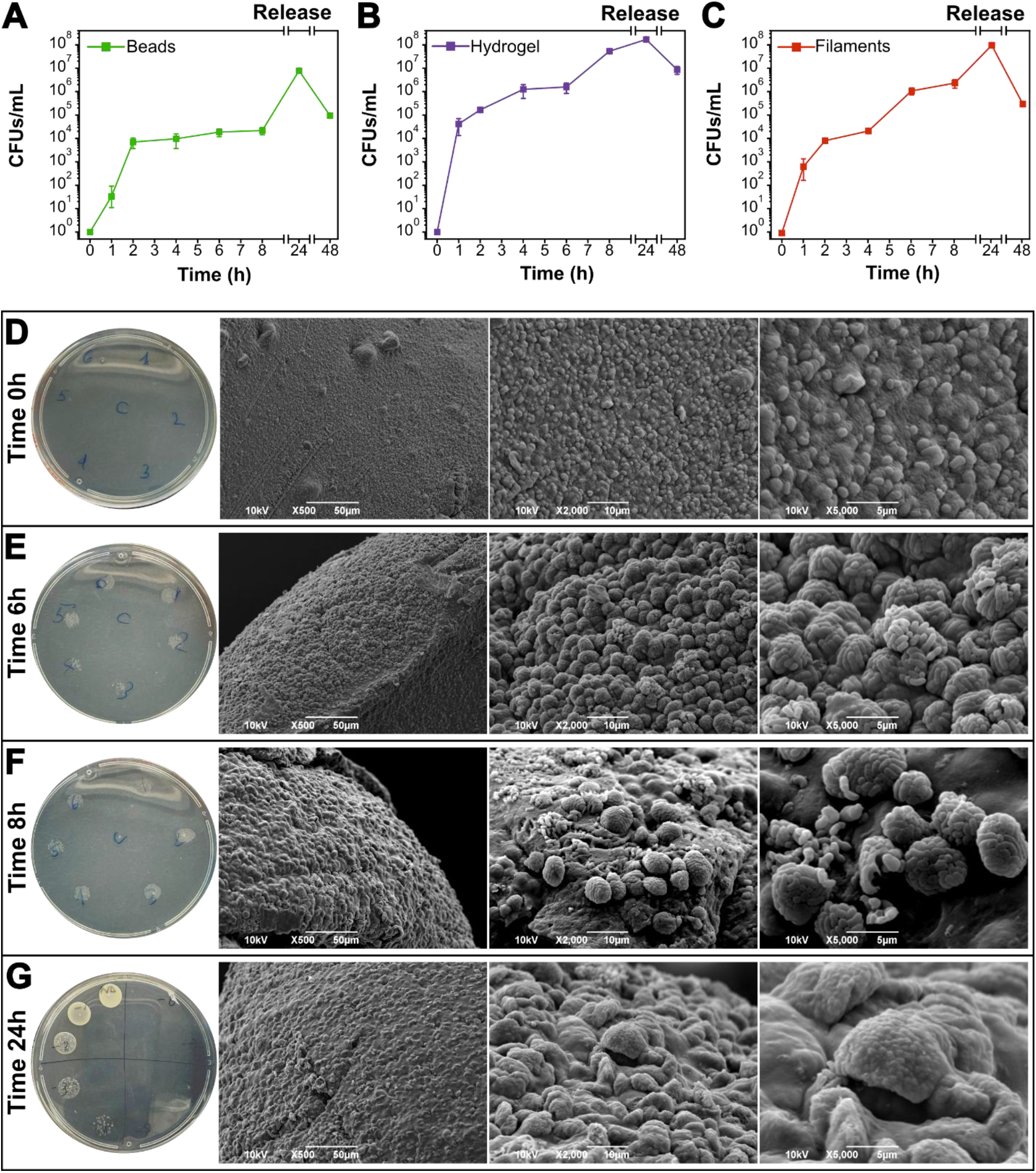
Probiotic test release *in vitro*. (A, B, and C) Probiotic release curve for beads, hydrogels, and filaments, respectively. (D, E, F, and G) Representative photographs of microbial agar plates and SEM images at different time points (0, 6, 8, and 24 hours) showing probiotic growth, the surface morphologies, and the presence of encapsulated probiotics on the beads during the release experiment.

For alginate beads, during the early incubation period, bacteria began to be released, and the concentration increased after 1 h, reaching 3.33 × 10¹ CFU mL⁻¹. After that, the bacterial amount was found to be 7.00 × 10³ CFU mL⁻¹ at 2 h, 9.66 × 10³ CFU mL⁻¹ at 4 h, 1.86 × 10⁴ CFU mL⁻¹ at 6 h, and 2.16 × 10^4^ CFU mL⁻¹ at 8 hours. The peak of the release was reached at 24 h (7.83 × 10⁶ CFU mL⁻¹), followed by a decline to 9.50 × 10⁴ CFU mL⁻¹ at 48 h, likely attributable to nutrient depletion.

A comparable release profile was observed for hydrogels and filaments, characterized by an initial exponential increase indicative of a burst release followed by a reduction in the release rate. For hydrogels, after 2 h, the concentration was 1.65 × 10⁵ CFU mL⁻¹, at 4 h, it was 1.25 × 10⁶ CFU mL⁻¹, at 6 h, it was 1.58 × 10⁶ CFU mL⁻¹, at 8 h, it was 5.33 × 10⁷ CFU mL⁻¹, and at 24 h, the maximum concentration was obtained (1.70 × 10⁸ CFU mL⁻¹). At 48 h, the concentration decreased to 8.66 × 10⁶ CFU mL⁻¹.

Filaments exhibited a similar trend, with a concentration of 6.66 × 10² CFU mL⁻¹ after 1 h, increasing to 8.50 × 10³ CFU mL⁻¹ at 2 h, 2.18 × 10⁴ CFU mL⁻¹ at 4 h, and 1.11 × 10⁶ CFU mL⁻¹ at 6 h. The concentration reached 2.40 × 10⁶ CFU mL⁻¹ at 8 h and peaked at 24 h (9.75 × 10⁷ CFU mL⁻¹). A subsequent decrease to 3.07 × 10⁵ CFU mL⁻¹ was observed at 48 h.

The release profiles of blank systems and bacterial solutions were studied as negative and positive controls, and the results are presented in **Figure S6**. No bacteria were observed in the blank systems, whereas the bacterial solutions showed a quick exponential growth followed by a plateau. These findings are further corroborated by CFUs observed on agar plates and by SEM images of beads, chosen as the representative model among the proposed systems, as shown in **Figures 3D, E, F**, and **G**, and **Figure S7**. Immediately after immersion in seawater, no CFUs were observed on agar plates. At the SEM, the rounded appearance of the probiotics on the material surface could be observed (**Figure 3D**), as previously described in **Figure 1H**, confirming that the probiotics were not released. In contrast, after 6 hours, the number of CFUs on agar plates was visibly higher and countable. In SEM images, the material surface appeared colonized by fully released probiotics, which were abundant, forming clusters and exhibiting the elongated shape observed for probiotics in the MB culture control (**Figure 3E**). These elongated bacterial clusters remained visible on the surface after 8 hours, indicating that the release had reached a stable rate. This was also confirmed by the number of CFUs present on agar plates (**Figure 3F**). After 24 hours, the probiotics are hardly visible at the SEM images of the surface, which show parts of the alginate material lifted, highlighting the probiotic escape from the structures (**Figure 3G**). Internal observation of the bead structure after 24 h uncovered the presence of a small number of bacteria still entrapped in the matrix (**Figure S8**). Nevertheless, the increasing CFUs on the agar plates over time indicate that the released probiotics remained alive and capable of growth and replication. SEM control images of the three blank systems are shown in **Figure S9**.

### 2.5 Biocompatibility and applicability tests on coral species

The biocompatibility of the delivery systems was assessed across multiple coral species, including *Pocillopora damicornis*, *Heliofungia fralinae*, and *Montipora foliosa*, representing diverse coral morphologies. All materials were administered via gentle deposition on corals, and continuous water movement was simulated in the tanks using air stones and surface aeration. We first tested *Pocillopora damicornis*, a representative branched coral (**Figure 4A**). Upon deployment, the beads were easily applied to the coral surface and remained securely attached, while the hydrogels adhered effectively to the branches and maintained their position throughout the experiment. Manual placement of filaments underwater was challenging due to the spatial constraints imposed by the branches; however, once positioned, the fibers remained lodged between the branches.

**Figure 4.**
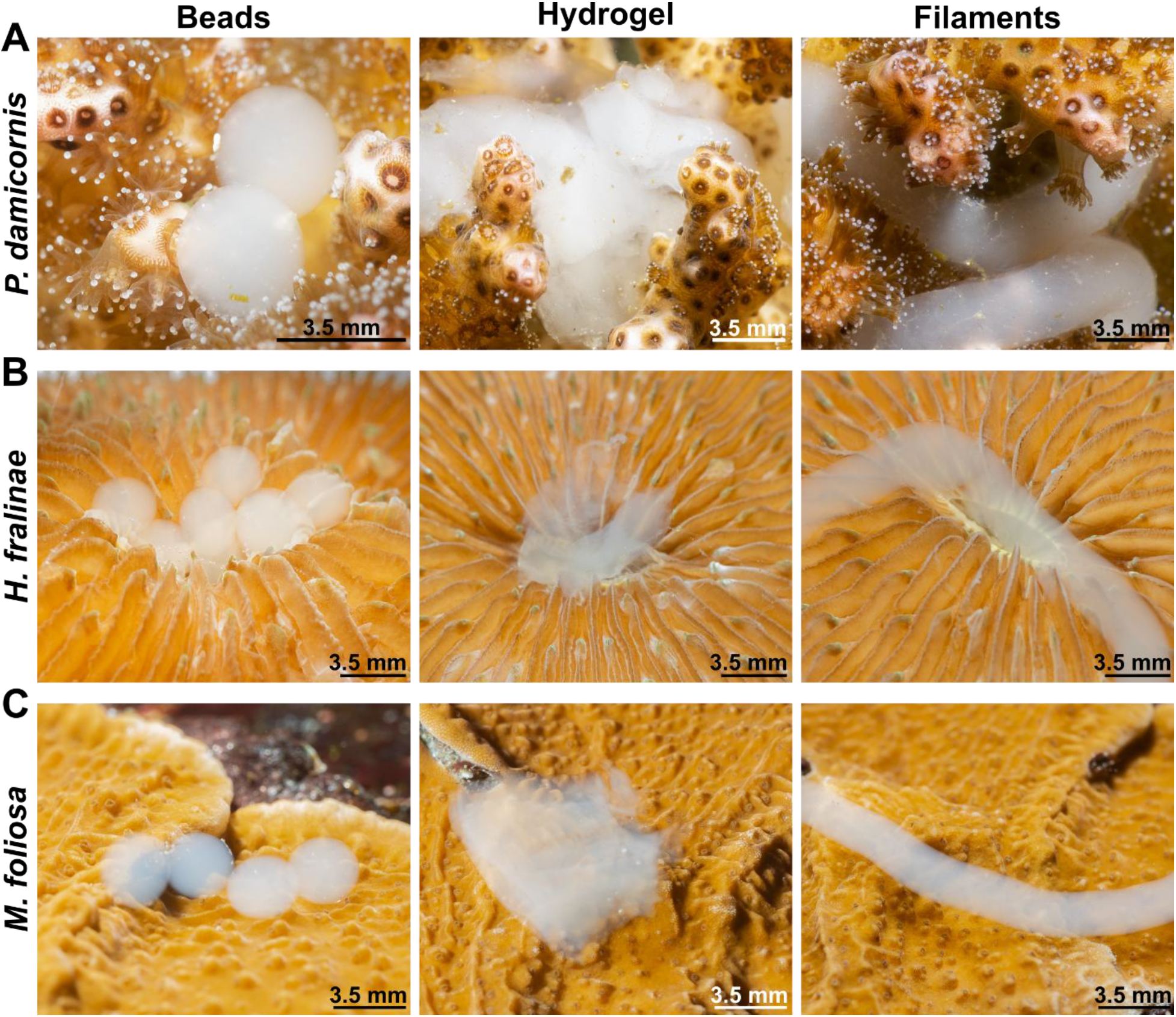
Application of probiotic delivery systems across coral species. (A, B, and C) Application of beads, hydrogel, and filaments on *Pocillopora damicornis*, *Heliofungia fralinae,* and *Montipora foliosa* corals, respectively.

On *Heliofungia fralinae*, a mushroom coral that possesses a large central oral opening, all three delivery systems remained stably attached within the coral mouth (**Figure 4B**). On *Montipora foliosa*, selected as a representative flat coral species due to its plate-like foliose growth, the beads and filaments adhered well, while the hydrogel showed slight instability under simulated water-flow conditions (**Figure 4C**). Across all species, the application of these probiotic delivery systems did not compromise coral health. Fully extended polyps and the absence of visible signs of stress, such as tissue necrosis, color changes, or excessive mucus production, indicated that the treatments were well tolerated and biocompatible.

### 2.6 Underwater probiotic delivery to corals

An *in vivo* delivery test was conducted on *Pocillopora damicornis* to evaluate the ability of probiotics to be released from different vehicles, reach corals, and integrate into the microbiome. The experimental design for the *in vivo* probiotic delivery test is shown in **Figure 5**. Five separate experimental tanks were used: the first three contained coral nubbins treated with beads, hydrogels, and filaments, respectively. The fourth had corals injected with a probiotic liquid culture using a syringe, while the last one served as an untreated control (**Figure 5A**). The experiment was conducted over a 72-hour period, with samples collected at 0, 8, 24, 48, and 72 hours (**Figure 5B**). Coral fragments were sampled at each time point, and microbiome analysis to assess probiotic delivery was performed (**Figure 5C**). A few details on the overall results of Illumina sequencing are reported in the Supplementary Materials.

**Figure 5.**
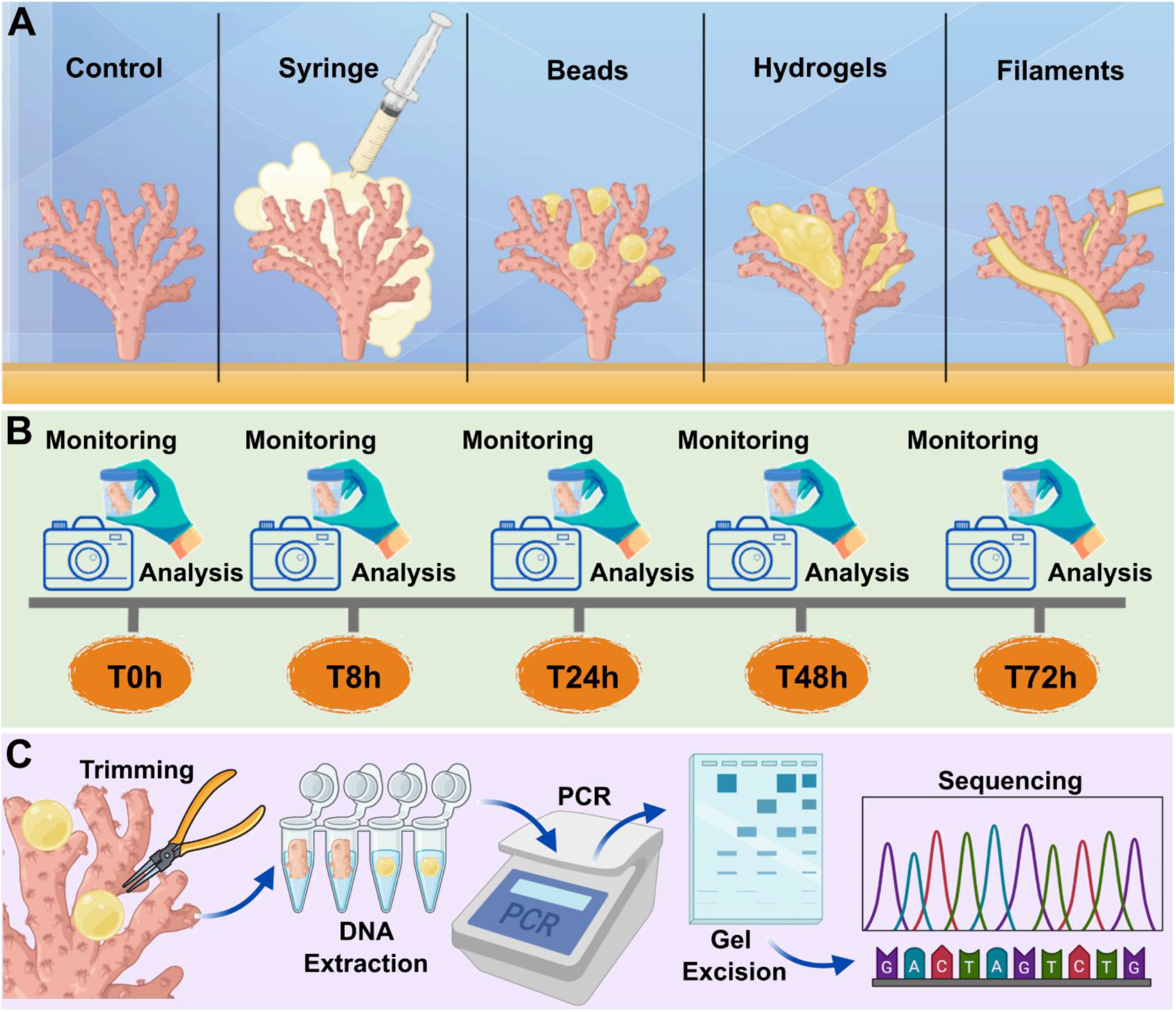
Schematic representation of the *in vivo* probiotic release experiment. (A) Experimental setup showing the division of *P. damicornis* coral fragments into five treatment groups: control, probiotic injection via syringe, and treatment with three delivery systems (beads, hydrogels, filaments). (B) Experimental timeline indicating coral monitoring over a 72-hour period with sampling at time points 0, 8, 24, 48, and 72 hours. (C) Overview of the microbiome analysis workflow, including coral tissue trimming, DNA extraction, PCR amplification, gel excision, and sequencing.

The relative abundance of sequences classified as genus *Pseudoalteromonas* is reported in **Figures 6A, B, C**, and **D**, and **Table S10**, while the results for all major families identified are provided in **Table S11**. In control corals (**Figure S10A**), the relative abundance of *Pseudoalteromonas* within the total microbiome remained at physiological baseline levels (1.98% ± 2.15%). Similarly, the syringe treatment (**Figure 6A**) did not induce substantial deviations from this baseline, with *Pseudoalteromonas* abundance remaining relatively stable throughout the experimental period. Specifically, the relative abundance after syringe injection was 0.75% ± 0.76% at 8 h, increased slightly to 1.57% ± 1.14% at 24 h, declined to 0.65% ± 0.69% at 48 h, and rose again to 1.47% ± 1.99% at 72 h. In contrast, corals treated with beads (**Figure 6B**) showed a marked, transient enrichment of *Pseudoalteromonas*. Relative abundance increased from 1.39% ± 2.62% at 8 h to 6.15% ± 2.99% at 24 h, reaching a maximum of 11.47% ± 10.69% at 48 h, before declining to 2.05% ± 1.67% at 72 h. During hydrogel treatment (**Figure 6C**), *Pseudoalteromonas* abundance remained comparatively stable between 8 h (9.33% ± 14.8%) and 24 h (8.30% ± 7.20%). A peak was observed at 48 h, with a relative abundance of 12.71% ± 5.88%, followed by a sharp decrease to 1.71% ± 2.41% at 72 h. The filament treatment (**Figure 6D**) also promoted a substantial increase in *Pseudoalteromonas* abundance over time. Relative abundance rose from 0.30% ± 0.41% at 8 h to 12.60% ± 12.40% at 24 h, decreased to 4.98% ± 7.41% at 48 h, and subsequently increased again to 9.38% ± 8.99% at 72 h. Two-sample t-tests revealed no statistically significant differences in the percentage of *Pseudoalteromonas* retrieved from corals subjected to different treatments or at different time points. However, in a few pairwise comparisons, the hydrogel treatment at 48 h or the bead treatment at 24 h could be considered only marginally non-significant (adjusted p-values ranging from 0.055 to 0.068; **Table S12**).

**Figure 6.**
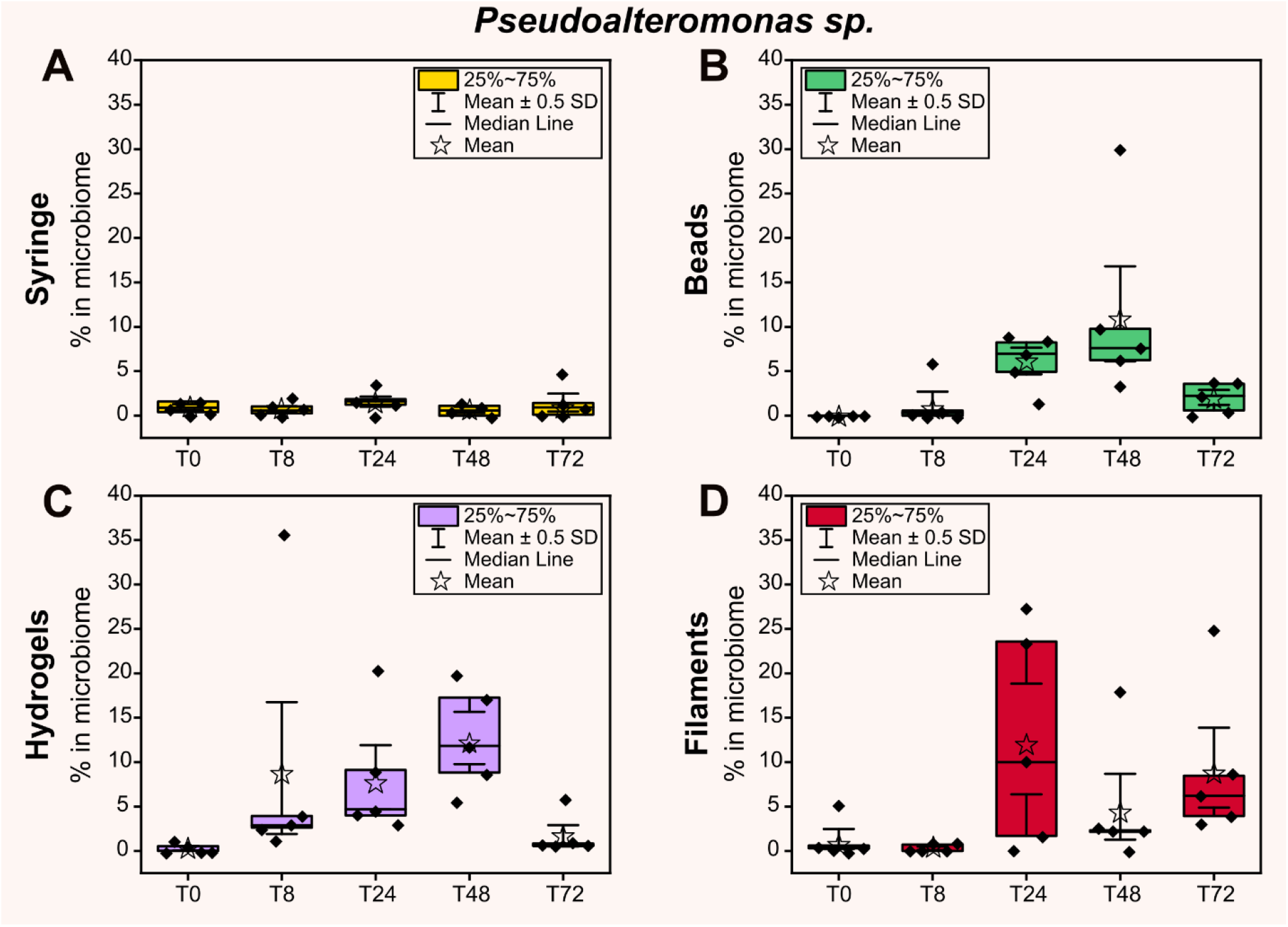
Relative abundance (%) of *Pseudoalteromonas* sp. within the coral microbiome during the four in vivo treatments. (A, B, C, and D) Relative abundance of *Pseudoalteromonas* spp. during the syringe, beads, hydrogels, and filaments treatments, respectively.

These results indicate that these delivery approaches were effective at temporarily increasing the relative abundance of *Pseudoalteromonas* in the coral microbiome. On the other hand, the application of the three alginate-based delivery systems was occasionally associated with a transient increase in the relative abundance of *Vibrio* spp. and members of the family *Cellvibrionaceae*. The results regarding these bacteria and the overall microbiome are reported in **Figures S10B**, **S10C**, **S11**, **S12**, and **S13.**

Redundancy analysis (RDA) was carried out to compare the prokaryotic community associated with corals across all experimental groups, and the results are shown in **Figure S14A**. Despite the similarities observed among coral microbiomes, the RDA model was overall significant. Post hoc tests generally revealed significant differences in the coral microbiome among the experimental groups (untreated control corals and corals treated with syringe, beads, hydrogel, and filaments). However, no significant differences were observed in the bacterial community between the beads and the controls (Padj = 0.118) or between the hydrogels and the filaments (P_adj_ = 0.079).

Finally, a schematic comparison of the results obtained in this work with previous attempts at probiotic inoculation reported in the literature is presented in **Figure S14B**. As can be observed, several inoculations are often required to achieve sufficient variation of the administered probiotic in the microbiome. On the other hand, our systems yielded the highest efficacy with the fewest inoculations (only one).

### 2.8 Potential mechanisms of breakdown and bioerosion of the probiotic delivery systems

The degradation of alginate-based delivery systems is generally driven by a combination of biological and chemical processes, enabling the controlled, site-specific release of probiotics. A schematic representation of these overlapping degradation mechanisms is shown in **Figure 7A**. To investigate the biodegradation mechanism, blank and loaded beads were placed underwater in contact with corals and analyzed after 72 hours for morphology and for the presence of microorganisms and ions. Morphological analysis of a blank bead is shown in **Figure 7B**, confirming the presence of microorganisms on its surface. Afterward, microbiome analyses of blank beads (**Figure 7C**) revealed that, at the beginning of the *in vivo* experiment, no bacterial taxa were detected, consistent with the sterile composition of the beads prior to coral exposure. However, after 72 h underwater, in contact with the coral, the beads showed substantial microbial colonization (and even after 20 min, see **Figure S15A**), characterized by a marked increase in *Pseudoalteromonas*, *Vibrio*, and members of the family Cellvibrionaceae, the latter of which are well-known polysaccharide degraders (*71*).

**Figure 7.**
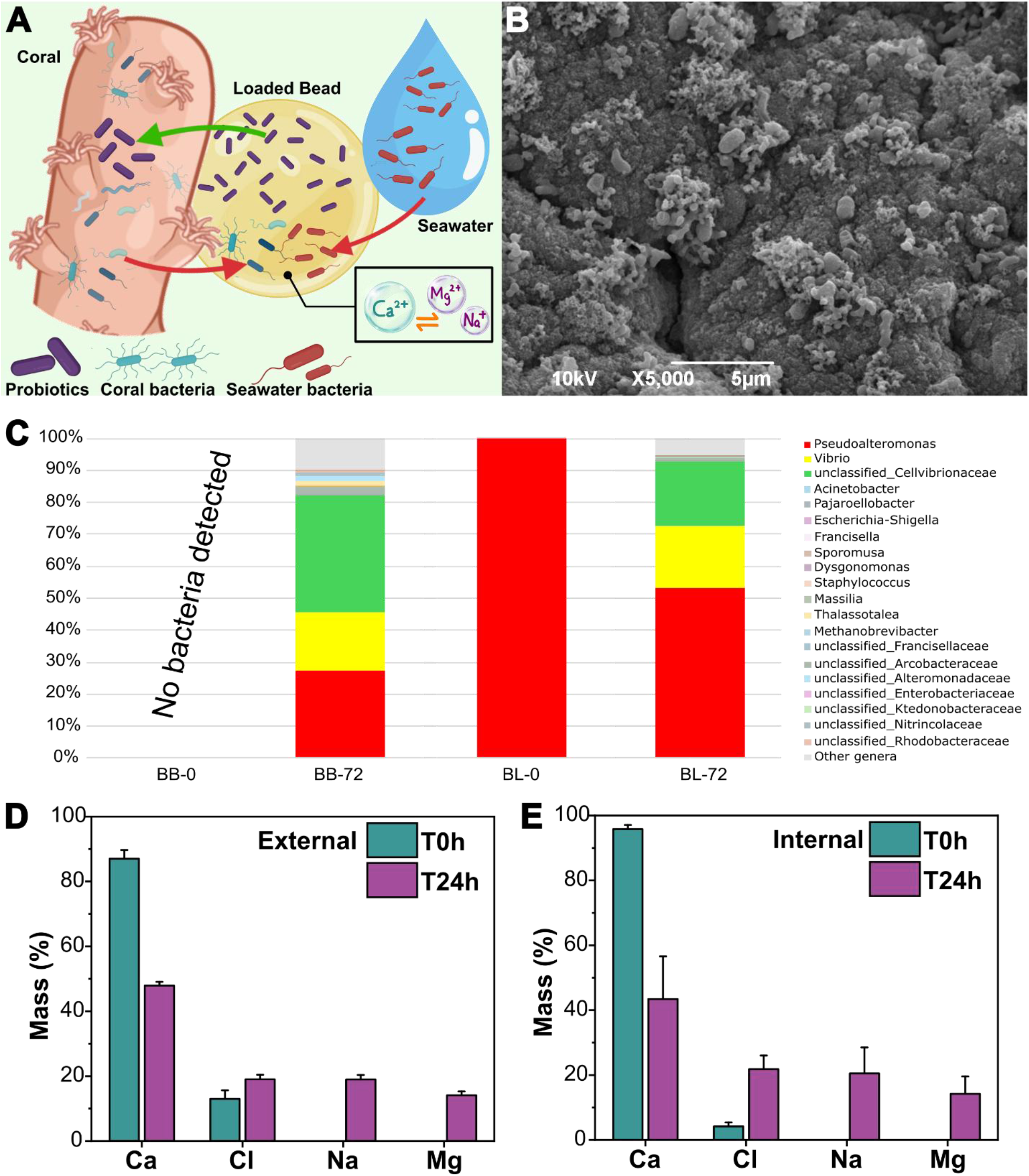
Degradation mechanism of the delivery systems. (A) Schematic illustration of the primary mechanisms involved in the degradation of the delivery systems. (B) SEM images showing colonization of the unloaded beads by various bacterial strains following exposure to coral for 72 h. (C) Microbiome results on blank and loaded beads at the beginning of the experiment and after 72 hours of immersion in seawater in contact with the corals. (D, E) EDX-SEM analysis of the surface and the cross-section of beads before and after being placed in seawater for 24 hours. Values of Ca, Cl, Na, and Mg are reported.

In contrast, beads loaded with the probiotic *Pseudoalteromonas* sp. were dominated by this genus, accounting for 99.9 ± 0.0% of the detected ASVs at the beginning of the experiment. After 72 h, the relative abundance of *Pseudoalteromonas* decreased to 53.4 ± 11.2%, concomitant with an enrichment of *Vibrio* and *Cellvibrionaceae*, which emerged as the predominant additional bacterial populations detected in the samples. Finally, RDA analysis (**Figure S14A**) revealed that the communities developed on alginate beads, both blank and loaded, were clearly distinct from control coral microbiomes (**Figures 7C** and **S15B**).

The presence of ions on the blank and loaded beads after seawater immersion was assessed by EDX-SEM, and the results are shown in **Figures 7D** and **E**. The investigation of the presented elements revealed that, prior to immersion in seawater, the surface of the beads contained 87.1% Ca²⁺ and 12.9% Cl⁻, whereas the internal region exhibited 94.4% Ca²⁺ and 5.6% Cl⁻. These two ions were used to cross-link the alginate ionically. After 24 h of immersion in seawater, the Ca²⁺ content decreased markedly, while Cl⁻ increased, and additional ions, such as Na⁺ and Mg²⁺, were detected in both regions. Specifically, at the bead surface, the Ca²⁺ mass percentage decreased to 47.9%, whereas Cl⁻, Na⁺, and Mg²⁺ accounted for 19.0%, 19.0%, and 14.1%, respectively. A similar trend was observed in the internal region, where the Ca²⁺ content declined to 43.4%, while Cl⁻, Na⁺, and Mg²⁺ increased to 21.9%, 20.5%, and 14.2%, respectively. These results confirm that ion exchange between divalent calcium ions and mono- and divalent ions present in seawater drives progressive destabilization of the alginate crosslinked network. Specifically, exchange between Ca²⁺ ions within the alginate matrix and Na⁺ and Mg²⁺ ions in seawater after 24 h leads to chemical weakening of ionic crosslinks, resulting in gradual matrix disintegration (*72*). The combined effects of internal and external microbial activity and ion exchange, as well as water-mediated swelling, can ensure a sustained, localized release of viable probiotics and the complete degradation of the delivery system over time.

## Discussion

Here, we unlocked the possibility of administering probiotics on corals via a new route using biodegradable natural vehicles to significantly enhance microbiome manipulation and, consequently, to improve therapeutic efficacy. Three delivery systems were developed as beads, hydrogels, and filaments made of calcium alginate and loaded with the probiotic *Pseudoalteromonas peptidolytica*. The localization of the probiotic within the biopolymer matrix slightly reduced the initial mechanical strength of the materials compared to the blank samples. This is because bacteria introduce soft inclusions and structural heterogeneities into the polymer network. Bacterial cells disrupt the continuity of the ionically crosslinked alginate matrix, locally reducing crosslink density and increasing porosity, thereby lowering the Young’s modulus (*73*). On the other hand, during the 7-day tests, bacteria increased the vehicles’ overall stability over time. This mechanical stability can likely be explained by the presence of probiotics within the alginate matrix, which may reduce water ingression into the polymer matrix via steric hindrance, or they may strongly interact with the polymer matrix, contributing to structural reinforcement and system stabilization during seawater exposure. Indeed, over time, the metabolically active probiotics encapsulated inside the material can produce extracellular polymeric substances (EPS) and form biofilms that progressively occupy the pore space and establish additional physical and chemical interactions with the surrounding network (*74, 75*).

The probiotic release profiles were governed by the concurrent contributions of bacterial diffusion, matrix swelling and erosion, and bacterial growth. Four main phases driven by different mechanisms can be described. Within the first one to two hours, an initial burst release is observed upon immersion of the materials in the aqueous medium, consistent with water-induced swelling and the consequent diffusion of bacteria into the surrounding medium. Notably, among the three systems, the hydrogel showed the fastest and strongest burst release within the first hour. This is followed by a steady-state release mediated by diffusion, in which swelling slows, and the bacteria released into the medium are in the initial lag phase, with a very small increase in the overall number of colonies observed (between 2 and 6 hours). Between 8 and 24 hours, a second increase in bacterial abundance is observed, which may result from progressive water-mediated erosion of the matrix and from bacterial growth within the material and in the release medium. Finally, after 24 hours, the bacterial concentration decreases, coinciding with the late stages of both the release and growth profiles. This decline is consistent with nutrient depletion in the medium, which ultimately limits bacterial proliferation and reduces the measured bacterial concentration. The beads showed well-defined separation of the 4 phases, whereas hydrogels and filaments exhibited greater overlap among these mechanisms.

The three developed delivery systems are versatile enough to be applied to a wide range of coral morphologies (branched, solitary, and flat), enhancing their applicability across diverse reef systems, as shown in **Figure 4**. In all cases, the loaded materials were “coral-compatible”, with no signs of toxic reaction. After the *in vivo* administration of *Pseudoalteromonas*, microbiome analyses over time were conducted. The natural abundance of this genus in the native coral microbiome ranged from 1–2%. Injecting a bacterial suspension into the water column with a syringe did not yield a measurable enrichment of probiotic bacteria in the coral microbiome, which remained comparable to that observed in untreated control corals. On the other hand, an increase in the relative abundance of *Pseudoalteromonas* was observed in corals treated with all three formulations, starting at 8 hours and peaking at 48 h. In particular, the beads and the hydrogels treatment at 48 h and the filaments treatment at 24 hours enabled the recovery of very high percentages of *Pseudoalteromonas* in coral microbiomes, reaching approximately 12%, 13%, and 13% on average, respectively, with the highest values of ∼20–30%, at least for some individuals (n = 7, out of 30 samples at 24 and 48 h), reaching peaks of concentrations nearly tenfold higher than the observed physiological levels. In contrast, the highest relative abundance of the probiotic, obtained via syringe injection, did not exceed 4.9% (observed in only 1 of 20 treated via this route across all time points), with values generally below 2%.

These observations provide strong evidence that the delivery strategies successfully promoted probiotic association with the coral host, thereby demonstrating a crucial role in potentially enhancing the probiotic therapy strategy. Despite this clear trend, the overall statistical support for the treatment effect was weakened by the exceptionally high variability observed among biological replicates (see **Table S12**). Indeed, within 24 - 48 hours after application of the three vehicles, 27 out of 30 fragments showed an increase in *Pseudoalteromonas* abundance relative to the physiological amount (above 2%), indicating a 90% success rate. Consequently, the high inter-sample variability reduced the statistical power necessary to detect consistent differences at the population level. Importantly, the lack of statistical significance should not be interpreted as evidence of treatment inefficacy, but rather as a reflection of the intrinsic heterogeneity of coral-associated microbial communities and the complex dynamics that govern host–microbe interactions *in vivo*.

Aside from the absence of adverse effects associated with the transient enrichment of potential pathogenic strains and the effective introduction of the probiotic, another key parameter for ensuring the suitability and ecological compatibility of a probiotic application is the overall composition of the host microbiome. In our experiment, as indicated by the RDA results, the communities on the coral treated with beads were comparable to and not significantly different from those on the untreated controls. This suggests that the vehicle can successfully deliver the probiotic without altering the overall microbiome and appears to be the least impactful treatment on corals, helping maintain the microbiome’s unaltered ecological roles. Hydrogels and filaments were found to perturb the coral microbiome relative to the control, but no differences were observed between them. However, they successfully delivered *Pseudoalteromonas.* Interestingly, syringe injection induces a shift in the coral microbiome while delivering probiotics with low efficiency, further confirming its overall unsuitability as a delivery method.

The efficacy of probiotic delivery by syringe injection was insufficient, and this would be further decreased in an open field, where rapid dilution of the probiotic in the seawater volume and wave- and current-correlated movements occur. Instead, by controlling the spatial and temporal release of BMCs through delivery systems, probiotics were maintained at the target site for prolonged periods, thereby reducing premature clearance, improving local bioavailability, and ultimately maximizing pharmacological effectiveness (*76, 77*). Finally, comparing the outcomes obtained in this study with those reported in the literature (see **Figure S14B**) places this work as the most effective to date in terms of the percentage of delivered bacteria in the microbiome after a single inoculation, where the concentration of the delivered probiotic was less than 5% in relative abundance after a single inoculation (*48–50*).

Regarding the degradation of the developed delivery systems in the marine environment, after 72 hours of immersion in seawater, the materials underwent microbial colonization, mostly mediated by *Vibrio*, *Pseudoalteromonas*, and unclassified members of the family Cellvibrionaceae. The family *Cellvibrionaceae* comprises a metabolically versatile group of marine heterotrophic bacteria, well recognized for their ability to produce various carbohydrate-active enzymes (CAZymes), including polysaccharide lyases that depolymerize marine biopolymers such as alginate (*71*). Similarly, *Pseudoalteromonas* can contribute to long term matrix degradation by producing alginate lyases that digest alginate to use it as a carbon and energy source (*80, 81*). Simultaneously, ionic exchange between divalent cations in the alginate network and seawater ions further destabilizes the network chemically. Overall, the gradual biodegradation is confirmed to be mediated by microbial, enzymatic, and chemical factors.

These first-generation prototypes of probiotic delivery systems highlight a step forward in the potential of combining materials science and pharmaceutical principles to advance marine biological applications, particularly by developing natural, biodegradable systems for coral restoration and conservation, as recently described in the literature (*82–91*).

Future implementation of these platforms should focus on identifying optimal BMC consortia tailored to specific environmental contexts and coral species, and investigate the uptake, retention, and persistence of introduced microorganisms within the coral host (*6, 92*), as well as control the proliferation of other bacterial strains. Their efficacy under natural reef conditions, including both low- and high-energy environments, and across a broader range of coral species, should be assessed. These easy-to-handle systems could be deployed by divers (**Figure 1**) or autonomous robotic platforms, supporting repeated, targeted probiotic delivery for reef conservation and climate resilience.

## Conclusions

In the face of escalating threats from climate change and human activities, coral reefs urgently require innovative and scalable conservation strategies. This study presents the first biomaterial-based underwater delivery systems, biodegradable alginate-based beads, hydrogels, and filaments, designed for the targeted, site-specific administration of coral probiotics. These systems maintain high probiotic viability, support controlled release, can be applied to various coral shapes, and degrade naturally in the marine environment, offering a practical, versatile solution for both laboratory- and field-based microbial interventions.

By accommodating diverse coral morphologies and enabling the encapsulation of tailored BMC consortia, these systems represent a significant advancement in the application of microbial therapies for reef restoration. However, further research is essential to refine the selection of probiotic assemblages, assess host colonization dynamics, and evaluate the long-term persistence and ecological impacts of such interventions. Understanding the duration and effectiveness of probiotic-induced benefits will determine whether single or repeated applications are required, thereby shaping future restoration protocols.

Ultimately, these delivery systems connect materials science and pharmaceutics with marine microbiology, providing a flexible and environmentally friendly method to administer coral probiotics. Their development represents a key advancement in applying microbiome-based strategies for reef conservation and improving our overall understanding of host–microbe relationships in marine ecosystems, and the eventual presence of a third player, “the underwater vehicle”, that could be fundamental to coral health and resilience.

## 5. Materials and methods

### 5.1 Chemicals, materials, and probiotic bacteria

Alginic acid sodium salt (or sodium alginate, called ALG) with a viscosity of 15 - 25 cps, calcium chloride (CaCl_2_, MW = 110,98 g/mol), trisodium citrate dihydrate (MW = 294,10 g/mol), MB powder, agar powder, Na cacodylate buffer, and hexamethyldisilazane (HMDS) were purchased from Sigma-Aldrich. Paraformaldehyde (PFA), glutaraldehyde (GA), and osmium tetroxide were obtained from Agar Scientific. Filtered seawater (FSW) was obtained by filtering the water from the Aquarium of Genova (Genoa, Italy) through a sterile membrane filter (Stericup® Quick Release Millipore Express® Plus 0.22 µm PES) to remove impurities. All water referred to as Milli-Q was purified with a Millipore Milli-Q system (18.2 MW.cm; Merck) prior to use. The probiotic bacterium of the species *Pseudoalteromonas peptidolytica* (DSM-14001) was purchased from the Leibniz Institut DSMZ.

### 5.2 Preparation of alginate beads, hydrogels, and filaments

To prepare a 5% (w/v) sodium alginate solution, alginate powder was gradually stirred into MB and autoclaved at 121 °C for 15 min. For bead preparation, the sterile alginate solution (5% ALG+MB) was added dropwise into a CaCl_2_ 0.1 M solution by using a 10 mL syringe with an 18-gauge needle. The beads formed were kept stirred in the CaCl_2_ solution for 5, 10, 20, and 30 min to allow complete hardening of the alginate. For hydrogel fabrication, the 5% ALG+MB solution was combined with CaCl_2_ 0.2 M in a 1:1 ratio, mixed, and left hardening for 5, 10, 20, and 30 min. Filaments were produced by extruding the 5% ALG+MB solution through a 10 mL syringe with the diameter of the exit hole of 2.30 mm (without a needle) into CaCl_2_ 0.1 M solution, which was soaked for 5, 10, 20, or 30 min. After polymerization, all three systems were rinsed in FSW and dried on paper to remove excess CaCl_2_.

Prior to all subsequent analyses, the diameters of beads and filaments were measured with a caliber, and the filaments were then cut to a length of 3 cm. Hydrogels were prepared by cutting them using a circular insert with an 8 mm diameter, and the thickness was measured with a caliper.

### 5.3 Culture conditions and growth curve

The species *Pseudoalteromonas peptidolytica* (DSM-14001) was cultured in MB at 30 °C for 24 hours with shaking at 180 rpm. Aliquots of the strain were stored in 15% glycerol at −80 °C for future use. The abundance of bacterial cells was determined using the plate count method and spectrophotometric analysis. The growth curves both in MB and 5%ALG+MB (**Figure S16**) monitored over 48 hours in three 250 mL flasks, with optical density (O.D.) at 600 nm measured using a Beckman Coulter DU 800 spectrophotometer. O.D. measurements taken at 0, 2, 6, 24, and 48 hours were directly correlated with the number of colony-forming units (CFUs) at each time point. CFU counts were determined by serial dilutions (10^-1^ to 10^-8^) of triplicate samples (n = 3) from each flask, which were plated in triplicate on Marine Agar (MA).

### 5.4 Encapsulation of probiotic bacteria

An aliquot of *Pseudoalteromonas peptidolytica* stored at –80 °C was thawed and incubated at 30 °C for 24 hours in MB. Meanwhile, the 5% ALG+MB solution was sterilized. After the incubation period, a flask containing 100 mL of sterilized 5% ALG+MB solution and 2 mL of liquid culture was prepared. The flask was then incubated at 30 °C for 24 hours at 200 rpm. Finally, beads, hydrogels, and filaments loaded with probiotics were prepared according to the protocol described in Section 5.2 and used immediately after production.

The encapsulation efficiency (EE) of probiotics was determined following the protocol of Li et al., 2009 (*62*), using *n* = 10 samples for each system (beads, hydrogels, and filaments). For each system, five freshly prepared samples were incubated in 2.5 mL of trisodium citrate dihydrate solution (0.06 M) for 45 min to disrupt the cross-linked alginate network and release the encapsulated probiotics. A control containing non-encapsulated cells in the same trisodium citrate dihydrate solution was included to confirm that the solution did not affect cell viability. After complete dissolution of the samples, viable bacterial counts were determined by the spread-plating method. The remaining five samples from each system were dried and weighed using an analytical balance (Kern ABT model 120-5DM) to determine their dry mass. The encapsulation efficiency of the three alginate-based delivery systems was then expressed as the number of viable cells normalized to the dry weight (CFU/g polymer), allowing comparison of the probiotic loading per unit mass of polymer.

### 5.5 Morphological characterization

The morphology of the obtained delivery systems, both blank and with probiotics, was analyzed by Scanning Electron Microscopy (SEM), using a variable-pressure JEOL JSM-6490LA (JEOL, Tokyo, Japan) microscope equipped with a tungsten thermionic electron source. The specimens were coated with a 20 nm-thick gold layer utilizing a Cressington 208HR sputter coater (Cressington, Watford, UK). A secondary electron detector was used for acquiring morphological information about the samples in high vacuum mode, with an acceleration voltage of 10 kV. Elemental analysis and mapping were achieved by energy-dispersive X-ray spectroscopy (EDX; JEOL, 10 mm^2^ Si:Li detector), operating at 15 kV.

The three blank systems and *Pseudoalteromonas peptidolytica* alone were observed as controls. Later, beads, hydrogels, and filaments loaded with the bacteria were analyzed at different time points (0, 6, 8, 24, and 48 h) to observe changes in the morphology during microbial growth. All the samples were fixed for 4 hours at room temperature in a solution of 1% PFA + 2% GA in 0.1 M Na cacodylate buffer (pH= 7.4). After the fixation, the samples were washed with 0.1 M Na cacodylate buffer (pH 7.4) 3 times for 5 min each, post-fixed in 1% osmium tetroxide in Milli-Q water for 1 hour and washed with Milli-Q water 2 times. The samples were subsequently dehydrated with a series of ascending EtOH solutions (70%, 90%, 96% and 3 times 100%) for 10 min each. Samples were incubated with 3:1, 1:1 and 1:3 ethanol: HMDS solutions, and 100% HMDS. Finally, the samples were left to dry overnight before the SEM analysis.

### 5.6 Mechanical properties

The mechanical properties of the three systems were characterized by nanoindentation with the Chiaro Nanoindenter (Optics11, Netherlands), working in displacement control. The system uses cantilever probes to apply controlled indentation to the samples while evaluating the reaction force from the cantilever deflection, measured by interferometry. Blank and loaded beads, hydrogels, and filaments were soaked in CaCl_2_ for different crosslinking times (5, 10, 20, and 30 min). Furthermore, to assess the temporal evolution of the mechanical properties of the filaments in seawater, samples subjected to 20 min of crosslinking were immersed in seawater maintained at 28 °C for seven days, and their mechanical properties were measured daily.

All samples were measured with a ≈4 N/m stiffness, ≈8 μm radius probe. Samples were tested while in liquid (filtered seawater), with a metallic weight holding them to the bottom of the Petri dish and preventing them from floating. Tests were conducted in displacement control; displacement was applied and removed at 5 μm/s. To calculate the Young’s modulus, indentation curves were fitted with two different models depending on the shape of the curve: when adhesion was limited, the classical Hertz model for contact between elastic blunt bodies was applied; when indentation was followed by pronounced adhesion, the Johnson-Kendall-Roberts (JKR) model, which adds the contribution of adhesion forces to the calculation of contact area, was applied. Typical curves of both types of behavior, Hertzian and adhesive, are shown in **Figures S3D**, and **E**.

The mechanical properties of alginate beads at the macroscale were determined using a standard compression test on a 3365 Instron dynamometer (Instron, Norwood, Massachusetts, USA) equipped with a 10 N load cell. Blank beads and beads loaded with probiotics were soaked in CaCl_2_ for 5, 10, 20, and 30 min (n=5 specimens per time point). The sample sizes were then measured with a caliper and tested at a strain rate of 0.016 s^-1^. After the measurements, the bead’s strength was extracted from the force/displacement curves, and data were analyzed by calculating the mean and standard deviation. Moreover, to evaluate changes in the mechanical properties of the beads over time in seawater, samples crosslinked for 20 min were immersed in seawater at 28 °C for 7 days, with measurements taken daily.

Rheological measurements of hydrogels were performed employing an Anton Paar Modular Compact Rheometer (MCR 302). To assess the effect of crosslinking duration, both blank and loaded hydrogels were immersed in CaCl₂ solutions for 5, 10, 20, and 30 min. To investigate the evolution of rheological properties during seawater exposure, hydrogels were immersed in seawater at 28 °C for 7 days to simulate marine conditions. The viscoelastic properties of the swollen hydrogels were analyzed daily throughout the immersion period. All rheological measurements were conducted at 28 °C.

Initially, amplitude sweep tests were conducted at an angular frequency of 10 rad/s and were essential for determining the linear viscoelastic region (the range over which the test can be carried out without destroying the sample structure). The shear strain range for the amplitude sweep tests was set to 0.01%-100%. After determining the linear viscoelastic region, an amplitude of 0.05% was selected for frequency sweep tests, which covered a range from 0.1 to 100 rad/s.

### 5.7 Probiotic release *in vitro*

Probiotic-loaded systems (n = 3 at each time point) were placed in a 24-well plate, with each well containing 1 bead, hydrogel, or filament and 1 mL of MB. The release kinetics were monitored for 48 hours. At specific time points (0, 1, 2, 4, 6, 8, 24, and 48 hours), the solution was collected from each well, and O.D. was measured using a spectrophotometer before being plated in triplicate on MA using serial dilutions ranging from 10^-1^ to 10^-8^ (at time points 0, 1, 2, 4, 6, 8, 24, and 48 hours). A positive control (C^+^) containing only probiotics, in the amount of 32 µL of liquid culture for the beads (81 CFUs/mL), 127 µL for the hydrogels (445 CFUs/mL), and 96 µL for the filaments (336 CFUs/mL) to take into account the differences in the materials geometries, and a negative control (C^−^) containing blank systems were also included for reference (**Figure S17**). Moreover, growth curves generated in MB and 5% alginate solution were used to establish the relationship between OD and CFU mL⁻¹ (**Figure S16**)

### 5.8 Maintenance and growth conditions of corals

#### 5.8.1 Acclimatization tank

The biocompatibility of the delivery systems was first evaluated in coral species spanning distinct morphological groups, including *Pocillopora damicornis*, *Fungia fungites*, and *Montipora foliosa*, (n = 5 per species). All coral species were kept in a 3100 L tank with photoperiod 12 h:12 h light:dark. Water quality parameters were maintained at pH 8.2 and a salinity of approximately 36 ppt. Ammonia and nitrite concentrations remained below 0.01 mg L⁻¹ and were monitored twice weekly.

For microbiome analysis, the reef-building coral *Pocillopora damicornis* was used as the test species. Experiments were performed with fragments (∼6−10 cm in length, with n = 5 for each experimental group) obtained from large mother colonies/genotypes raised inside tanks of the Aquarium of Genoa. The nubbins were randomly divided among the different experimental groups. Before the experiment, fragments were kept under controlled conditions in a 3100 L acclimatization tank (photoperiod 12 h:12 h light:dark) for 1 month to recover. Each night, 50 L of freshwater containing 500 g of calcium hydroxide were added dropwise to the system to promote coral skeletal calcification. Colonies were fed twice a week with a food mixture in a solution containing *Tetraselmis suecica* algae and Rotifera to improve recovery. During the day, the tank was set up as a semi-open system, constantly supplied (about 300 L per hour) with previously filtered seawater (via UV filtration) pumped from a depth of 50 m outside the Foranea Dam of Genoa. The temperature was kept at a constant 25 °C. Corals were illuminated with an HQI lamp (400w 10,000K Nepturion BLV) at an average irradiance of 250 μmol photons m^-2^ s^-1^. At night, the tank was set up as a closed system, with the water circulation pump (Argonaut AV150-2DN-SB 220v) set to 10-13 m^3^ per hour to ensure a complete water change every 25 min. The water from the tank, once pumped, passed through a filtration system consisting of a sand filter (0.4 mm, Astralpool ARTIC) and a UV filter (Panaque 750 s, with four 40 W lamps embedded); the water was then returned to the tank.

#### 5.8.2 Experimental tanks

After acclimatization, *P. damicornis* fragments were transferred to five experimental tanks, each set to the same parameters. Specifically, during the day, the tanks (50 L) were set up as an open system that constantly received filtered seawater from another water circulation system within the Aquarium of Genoa, supplied with 40 L per hour. Before entering the experimental tanks, the water passes through a filtration system consisting of a sand filter (0.4 mm, Astralpool ARTIC) that removes coarse particles, through an ultraviolet filter (WEDECO KATADYN AG) to sterilize the water, and finally through a protein skimmer (N Sguassero, Italy) that removes the organic agglomerates. Corals (photoperiod 12 h:12 h light:dark) were illuminated by two LED X 200W 10000K at an irradiance of 250 μmol photons m^-2^ s ^-1^. Furthermore, the tanks were supplied with 2 porous air stones for aquaria to keep the water constantly oxygenated and in motion, and with a flow pump (EHEIM 2275-02-0, max flow: 1250 L/h) to have further biological filtration. Two aquarium heaters (NEWA Therm Eco 300w) gradually increased the experimental temperature from 25 °C to 28 °C at a rate of 1 °C every day to allow corals to acclimate. The temperature was maintained at 28 °C for the subsequent experiments. The water tank’s chemical-physical parameters (pH, nitrates, nitrites, ammonium, phosphates, salinity) were monitored twice a week.

### 5.9 Application of materials on corals and sampling

Five different tanks were used for this experiment. In the first three, beads, hydrogels, and filaments were applied on the nubbins of *P. damicornis* and left for 72 hours to ensure proper delivery. Beads were injected into the branches of the coral fragment using a Pasteur pipette, while hydrogels were injected using a 2.5 mL syringe that had been modified by cutting the hub to match the diameter of the barrel (8.9 mm). The filaments were manually wrapped around the nubbins. In the fourth tank, a probiotic liquid culture was injected into the corals using a 30 mL syringe as described in the literature (*40–42, 50*). The last tank served as a control, with no probiotics introduced.

A total of 110 coral fragments were collected for microbiome analyses from each treatment (control, syringe, beads, hydrogels, and filaments) at different time points (0, 8, 24, 48, and 72 hours), except for controls, which were collected at 0 and 72 h only. All treatments were sampled in five replicates. Specifically, to confirm successful probiotic delivery, coral fragments (∼1 cm in length) that were in contact with the delivery systems were removed and stored in 2 mL tubes containing RNA-later at −20 °C for microbiome analysis. Additionally, coral fragments subjected to syringe-injection treatment were collected at each time point and stored in the same manner. Moreover, to check the presence and persistence of the probiotic, control bead delivery systems, both blank and loaded with the bacterial strain, were sampled in quadruplicate at the beginning of the experiment and after 72 hours of contact with the coral, as well as freshly prepared beads stored before entering the tank system, for a total of 20 bead samples. These samples were also preserved in 2 mL tubes containing RNA-later and stored at −20 °C for subsequent microbiome analysis.

### 5.10 Microbiome analysis

Total DNA was extracted from approximately 0.025 g of ground coral or from a single bead using the DNeasy® Blood & Tissue Kit (Qiagen) according to the manufacturer’s instructions. A first PCR amplification was performed on the V1-V3 hypervariable regions of 16S rRNA gene for each sample to evaluate its quality on the undiluted template and on the 1:10 and 1:100 dilutions to identify possible inhibition or insufficient samples. A second PCR was then performed on the V4 hypervariable region of the 16S rRNA gene for each sample using OneTaq® DNA Polymerase (New England Biolabs, Massachusetts, USA) and 0.2 µM of each primer, in a final volume of 100 µL. Illumina adapters and 6-bp barcodes were added at the 5’ end of each primer. 515F and 806R primers, modified to match both Bacteria and Archaea, were used (*93–95*). The cycling conditions were: initial denaturation at 94 °C for 3 min; 28 cycles at 94 °C for 45 s, 50 °C for 45 s, and 72 °C for 45 s, and a final extension at 72 °C for 5 min. Amplicons were run on 2% agarose gels, purified with Wizard® SV Gel and PCR Clean-up System (Promega Corporation, Madison, WI), and quantified with Qubit® (Life Technologies, Carlsbad, CA). Amplicon libraries were prepared with the addition of standard Nextera indexes (Illumina, Inc., San Diego, CA) and sequenced with MiSeq Illumina platform (Illumina, Inc., San Diego, CA), using a 2 × 300 bp paired-end protocol. Amplicon Sequence Variants (ASVs) were inferred using the DADA2 pipeline (*96*), as described in (*97*). After taxonomic classification using the latest version of the SILVA database (Ref NR 99 v. 138.2), ASVs unclassified at the domain level were discarded, as they generally corresponded to coral mitochondria or protein-coding genes.

The structure of prokaryotic communities hosted by beads and corals was compared among treatments and at different experimental times using redundancy analysis (RDA) of Hellinger-transformed ASV abundance, as defined by 16S rRNA gene amplicon sequencing data. RDA was followed by post hoc pairwise comparisons, the significance of which was adjusted using the False Discovery Rate (FDR) method. Analyses were performed in R 4.4.1. Two-sample (unpaired) t-tests were used to compare the percentage of the delivered *Pseudoalteromonas* strain among pairs of treatments and/or time points. The obtained p-values were corrected for multiple comparisons using the Benjamini–Yekutieli false discovery rate (FDR) method, considering differences significant at adjusted p < 0.05 (*98*).

## Data availability

Raw sequences from Illumina sequencing were submitted to Sequence Read Archive (SRA-NCBI), under the BioProject ID PRJNA1489437.

## Acknowledgments

The authors would like to declare that part of this research was funded under the National Recovery and Resilience Plan, Mission 4, Component 2, Investment 1.4—call for tender no. 3138 of 16 December 2021, rectified by Decree no. 3175 of 18 December 2021 of the Italian Ministry of University and Research, and funded by the European Union NextGenerationEU. Project code: CN_00000033; Concession Decree no. 1034 of 17 June 2022, adopted by the Italian Ministry of University and Research; CUP: J33C22001200001; project title ‘National Biodiversity Future Center (NBFC)’. This work is part of the ‘Technologies for Sustainability’ flagship program of the Italian Institute of Technology.

We would also like to acknowledge BioRender (http://biorender.com/), which was used to produce Figures 1, 5, and 7 of the main manuscript.

