## Supplementary Files for "Unlocking Targeted Underwater Delivery of Coral Probiotics"

§ Co-last authors

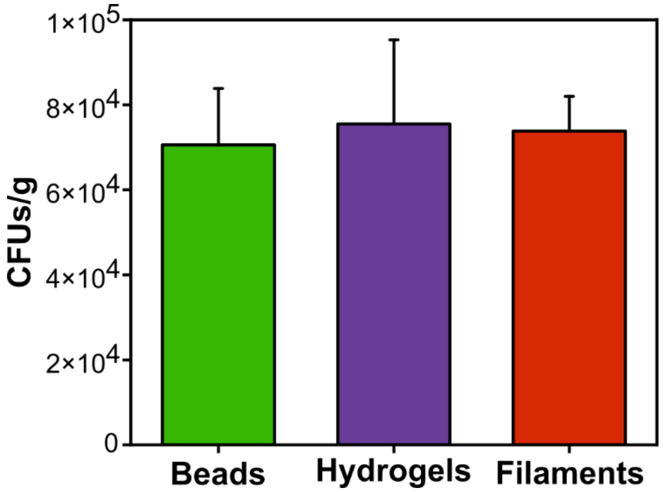

**Figure S1. Encapsulation efficiency.** Encapsulation efficiency for beads, hydrogels, and filaments, respectively.

| MEASURE | BEADS | HYDROGELS | FILAMENTS |
|---|---|---|---|
| | 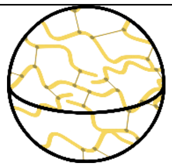 | 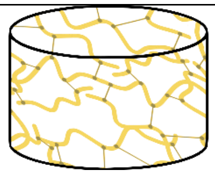 | 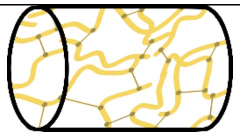 |
| Average wet mass of polymer (g) | 0.030 ± 0.001 | 0.342 ± 0.033 | 0.190 ± 0.035 |
| Average wet volume ($cm^3$) | 0.023 ± 0.004 | 0.319 ± 0.032 | 0.084 ± 0.021 |
| Average density ($g/cm^3$) | 1.330 ± 0.237 | 1.076 ± 0.108 | 2.309 ± 0.325 |
| Mean CFUs | 175 | 2875 | 2250 |
| Average dry mass of polymer (g) | 0.002 ± 0.0005 | 0.039 ± 0.009 | 0.030 ± 0.003 |
| Normalization CFUs/g polymer | 7.06E+04 ± 1326 | 7.55E+04 ± 1979 | 7.38E+04 ± 8136 |

**Table S1. Dimension measurements of the three delivery systems and encapsulation efficiencies.**

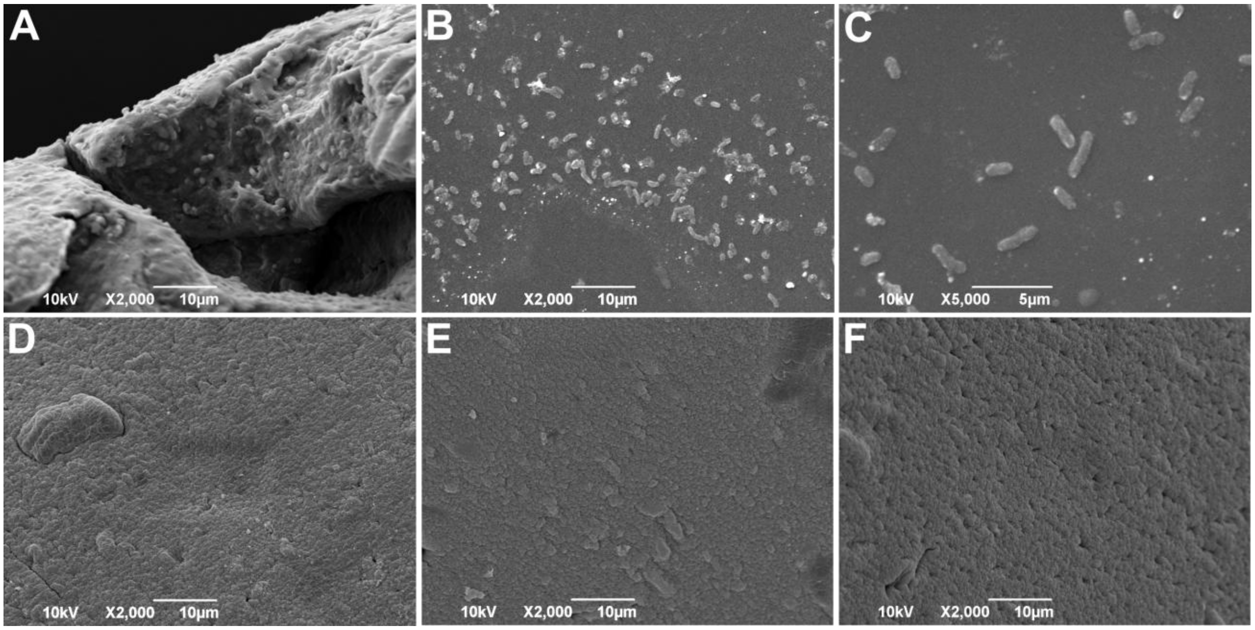

**Figure S2. SEM images of *Pseudoalteromonas peptidolytica*, beads, hydrogels, and filaments.** (A) Cross-section SEM image of a loaded bead confirming encapsulation of probiotics inside the polymer matrix. (B, C) Control SEM images showing the probiotic *Pseudoalteromonas peptidolytica* at two different magnifications. Here, their typical rod shape is clearly visible. (D, E, and F). Top-view SEM images showing blank beads, hydrogels, and filaments, respectively. All three blank systems exhibited smooth surfaces, consistent with the absence of probiotics.

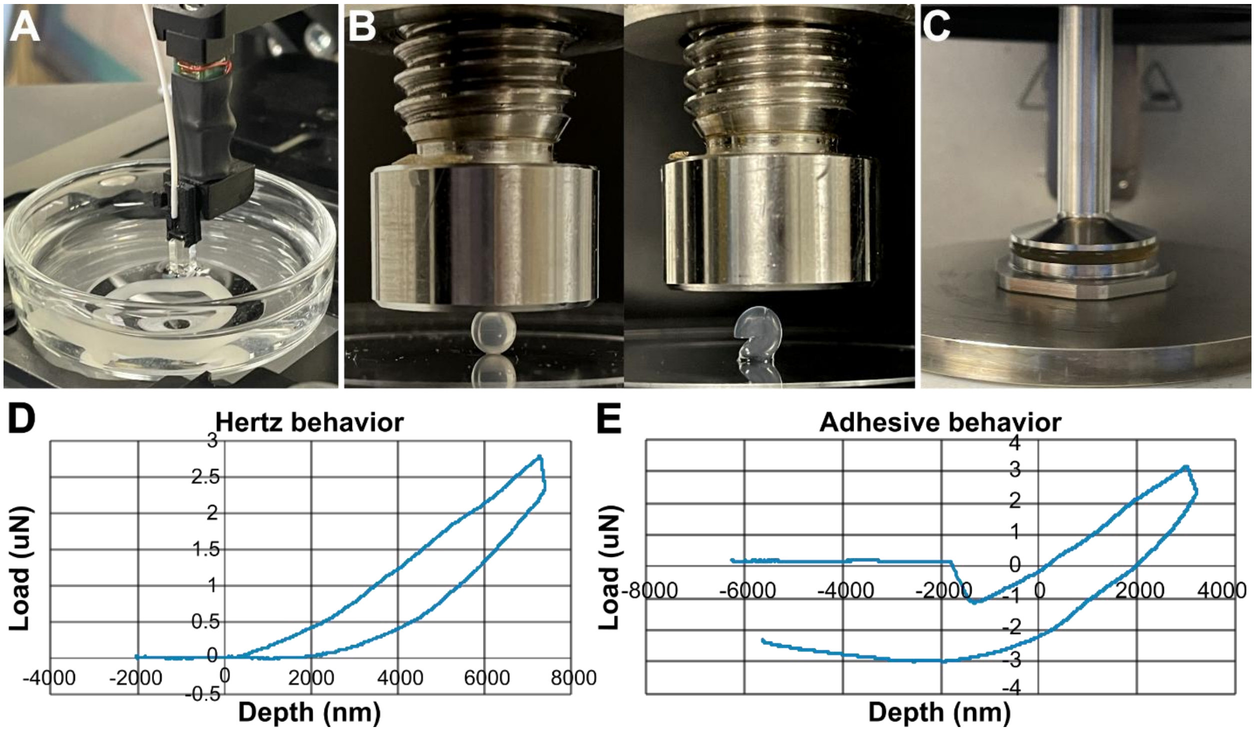

**Figure S3. Additional mechanical characterization of the material**. (A, B, and C) Photographs of nanoindentation, compression test, and rheometer. (D) Hertz fitting model for limited adhesion used to calculate Young's modulus. (E) Adhesive fitting model used to calculate Young's modulus.

| Beads | Blank | | Loaded | |
|---|---|---|---|---|
| Crosslinking Time | Young's Modulus E | Standard Deviation | Young's Modulus E | Standard Deviation |
| min | kPa | kPa | kPa | kPa |
| 5 | 56.787 | 5.704 | 54.115 | 4.331 |
| 10 | 58.370 | 8.419 | 67.552 | 6.129 |
| 20 | 74.768 | 6.457 | 61.884 | 6.358 |
| 30 | 61.163 | 2.676 | 62.859 | 4.228 |
| Hydrogels | Blank | | Loaded | |
| Crosslinking Time | Young's Modulus E | Standard Deviation | Young's Modulus E | Standard Deviation |
| min | kPa | kPa | kPa | kPa |
| 5 | 30.920 | 8.685 | 21.972 | 2.735 |
| 10 | 48.225 | 18.159 | 37.391 | 8.859 |
| 20 | 56.151 | 7.767 | 39.713 | 14.667 |
| 30 | 53.315 | 5.971 | 21.777 | 8.043 |
| Filaments | Blank | | Loaded | |
| Crosslinking Time | Young's Modulus E | Standard Deviation | Young's Modulus E | Standard Deviation |
| min | kPa | kPa | kPa | kPa |
| 5 | 48006.2215 | 5779.70422 | 46164.018 | 3229.31996 |
| 10 | 50347.65 | 5621.63665 | 47000.91393 | 2896.4379 |
| 20 | 53531.234 | 7583.8947 | 49235.35294 | 2666.09662 |
| 30 | 53710.7 | 10662.18771 | 51498.15442 | 1728.03933 |

**Table S2. Nanoindentation at different crosslinking times for beads, hydrogels, and filaments.**

| Filaments | Blank | | Loaded | |
|---|---|---|---|---|
| Day | Young's Modulus E | Standard Deviation | Young's Modulus E | Standard Deviation |
| | kPa | kPa | kPa | kPa |
| 0 | 53482 | 6595.79799 | 52132.30769 | 9068.41786 |
| 1 | 38542 | 1894.47618 | 50532.5 | 13280.11069 |
| 3 | 11270 | 2189.7387 | 50001.11111 | 3841.3391 |
| 5 | 3664.70588 | 1666.385 | 48113 | 7323.79084 |
| 7 | 0 | 0 | 41840 | 5641.27645 |

**Table S3. Nanoindentation of filaments immersed in seawater for 7 days.**

| Beads | Blank | | Loaded | |
|---|---|---|---|---|
| Crosslinking time | Maximum Force | Standard deviation | Maximum Force | Standard deviation |
| min | N | N | N | N |
| 5 | 0.698 | 0.093 | 0.700 | 0.014 |
| 10 | 0.742 | 0.095 | 0.724 | 0.032 |
| 20 | 0.883 | 0.036 | 0.786 | 0.023 |
| 30 | 1.231 | 0.115 | 0.830 | 0.037 |

**Table S4. Compression test on beads at different crosslinking times.**

| Beads | Blank | | Loaded | |
|---|---|---|---|---|
| Day | Maximum Force | Standard deviation | Maximum Force | Standard deviation |
| | N | N | N | N |
| 0 | 0.822 | 0.075 | 0.705 | 0.141 |
| 1 | 0.680 | 0.083 | 0.702 | 0.121 |
| 3 | 0.339 | 0.067 | 0.696 | 0.112 |
| 5 | 0.027 | 0.008 | 0.622 | 0.054 |
| 7 | 0.008 | 0.002 | 0.582 | 0.032 |

**Table S5. Compression test on beads immersed in seawater for 7 days.**

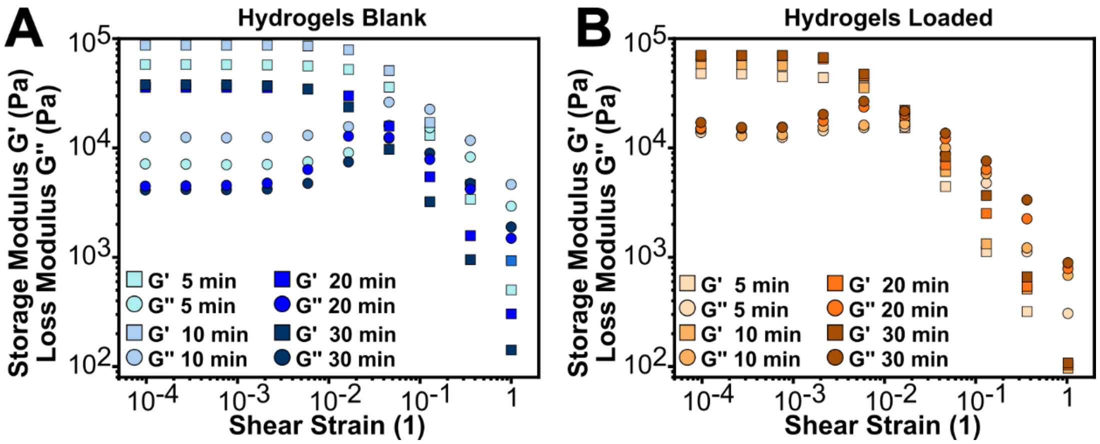

**Figure S4. Additional rheological characterization of the materials.** (A, B) Results of the amplitude sweep test performed on blank and loaded hydrogels at different crosslinking times.

| Amplitude sweep test | Blank | | | | | | | |
|---|---|---|---|---|---|---|---|---|
| Shear Strain | G' | G" | G' | G" | G' | G" | G' | G" |
| 1 | Pa | Pa | Pa | Pa | Pa | Pa | Pa | Pa |
| | 5min | 5min | 10min | 10min | 20min | 20min | 30 min | 30 min |
| 1.01E-04 | 57500 | 7103.5 | 86161 | 12479 | 35614 | 4115.1 | 37559 | 4455.5 |
| 2.80E-04 | 57499 | 7065.5 | 86296 | 12407 | 35582 | 4154.7 | 37594 | 4485.1 |
| 7.78E-04 | 57247 | 6987 | 86183 | 12318 | 35575 | 4128.4 | 37528 | 4531.5 |
| 0.00217 | 56921 | 7032.9 | 86040 | 12449 | 35382 | 4188 | 36977 | 4748.6 |
| 0.00603 | 55762 | 7441.6 | 84677 | 12966 | 34362 | 4725 | 34267 | 6326.9 |
| 0.0168 | 51800 | 9002.1 | 78375 | 15562 | 29791 | 7421.9 | 23566 | 12702 |
| 0.0466 | 35713 | 15941 | 50701 | 26035 | 15747 | 12212 | 9682.5 | 12417 |
| 0.13 | 12985 | 15202 | 16960 | 22425 | 5435.1 | 8907.4 | 3214.5 | 7821.3 |
| 0.361 | 3384.2 | 8219.2 | 4612.1 | 11650 | 1575.5 | 4720.7 | 952.37 | 4190.5 |
| 1.01 | 503.67 | 2926.2 | 933.72 | 4613.5 | 305 | 1890.3 | 142.81 | 1488.9 |

| Amplitude sweep test | Loaded | | | | | | | |
|---|---|---|---|---|---|---|---|---|
| Shear Strain | G' | G" | G' | G" | G' | G" | G' | G" |
| 1 | Pa | Pa | Pa | Pa | Pa | Pa | Pa | Pa |
| | 5min | 5min | 10min | 10min | 20min | 20min | 30 min | 30 min |
| 1.01E-04 | 44671 | 12967 | 54753 | 13827 | 63894 | 14211 | 65360 | 15964 |
| 2.80E-04 | 44335 | 12236 | 53759 | 12113 | 64819 | 14143 | 65116 | 14398 |
| 7.78E-04 | 41614 | 11690 | 52636 | 12360 | 64826 | 14481 | 65108 | 14421 |
| 0.00217 | 41005 | 13479 | 507291 | 14646 | 60782 | 16525 | 62314 | 18971 |
| 0.00603 | 34350 | 14347 | 33144 | 15081 | 42683 | 22089 | 44149 | 24873 |
| 0.0168 | 14342 | 14572 | 16367 | 15498 | 18554 | 19178 | 20694 | 20315 |
| 0.0466 | 4188.5 | 8254.5 | 5732.5 | 9434 | 6618.5 | 11407 | 7836 | 12772 |
| 0.13 | 1067 | 4491 | 1264 | 5462.5 | 2382 | 5970.5 | 3473 | 7131 |
| 0.361 | 305.24 | 1063 | 489.62 | 1155.9 | 517.66 | 2125.9 | 630.5 | 3159.6 |
| 1.01 | 50.367 | 292.62 | 92.997 | 650.89 | 99.664 | 749.16 | 103.85 | 846.98 |

**Table S6. Amplitude sweep test on blank and loaded hydrogels at different crosslinking times.**

| Frequency sweep test | Blank | | | | | | | |
|---|---|---|---|---|---|---|---|---|
| Angular Frequency | G' | G" | G' | G" | G' | G" | G' | G" |
| [rad/s] | Pa | Pa | Pa | Pa | Pa | Pa | Pa | Pa |
| | 5 min | 5 min | 10min | 10min | 20min | 20min | 30min | 30min |
| 628 | 56255 | 9982 | 59617 | 10883.9 | 62621 | 12981 | 65566 | 15036.4 |
| 628 | 49513 | 9194.1 | 53463 | 10478.2 | 56502 | 12176 | 59502 | 15164.4 |
| 628 | 45536 | 8665.6 | 47039 | 10422.3 | 50989 | 11662 | 53793 | 14718.7 |
| 215 | 40194.5 | 8685.2 | 42732.3 | 9946.9 | 45897 | 11598 | 48344 | 14484.1 |
| 59.9 | 35123 | 8339.8 | 38934.7 | 10133.5 | 41036 | 11924 | 44405 | 14353.2 |

| | | | | | | | | |
|---|---|---|---|---|---|---|---|---|
| 16.7 | 29628.2 | 8991.4 | 33249.9 | 11076 | 36021 | 12549 | 39594 | 14873.1 |
| 4.64 | 27457.7 | 9719.02 | 28617 | 11550.2 | 31956 | 13055 | 34840 | 15229.1 |
| 1.29 | 20372.6 | 9555.02 | 24041.4 | 11644.9 | 27812 | 13212 | 30175 | 15709 |
| 0.359 | 19328.1 | 9452.53 | 21545.4 | 11008.2 | 24625 | 13169 | 27589 | 15177.8 |
| 0.1 | 17316.2 | 9398.51 | 19562 | 11114.9 | 22881 | 13651 | 25013 | 15212.1 |
| Frequency sweep test | Loaded | | | | | | | |
| Angular Frequency | G' | G" | G' | G" | G' | G" | G' | G" |
| [rad/s] | Pa | Pa | Pa | Pa | Pa | Pa | Pa | Pa |
| | 5 min | 5 min | 10min | 10min | 20min | 20min | 30min | 30min |
| 628 | 17822 | 2941 | 22496 | 3911 | 28479 | 6438 | 31976 | 8091 |
| 628 | 16706.5 | 2862 | 21380.5 | 3832 | 27364 | 6359 | 30860 | 8012 |
| 628 | 16065 | 2953.5 | 20739 | 3923.5 | 26722 | 6451 | 30219 | 8103.5 |
| 215 | 14450 | 1865.5 | 19124 | 2835.5 | 25107 | 5363 | 28604 | 7015.5 |
| 59.9 | 12474 | 905 | 17148 | 1875 | 23131 | 4402 | 26628 | 6055 |
| 16.7 | 13853 | 917 | 18527 | 1887 | 24510 | 4414 | 28007 | 6067 |
| 4.64 | 12370.5 | 1468 | 17044.5 | 2438 | 23028 | 4965 | 26524 | 6618 |
| 1.29 | 10946 | 1450 | 15620 | 2420 | 21603 | 4947 | 25100 | 6600 |
| 0.359 | 9535 | 2194 | 14209 | 3164 | 20192 | 5691 | 23689 | 7344 |
| 0.1 | 9119 | 2035.5 | 13793 | 3005.5 | 19776 | 5533 | 23273 | 7185.5 |

**Table S7. Frequency sweep test on blank and loaded hydrogels at different crosslinking times.**

| Amplitude sweep test | Blank | | | | | | | | | |
|---|---|---|---|---|---|---|---|---|---|---|
| Shear Strain | G' | G" | G' | G" | G' | G" | G' | G" | G' | G" |
| 1 | Pa | Pa | Pa | Pa | Pa | Pa | Pa | Pa | Pa | Pa |
| | T0 | T0 | T1 | T1 | T3 | T3 | T5 | T5 | T7 | T7 |
| 1.00E-04 | 54468 | 12837 | 19480 | 3286 | 28753 | 4517.2 | 15649 | 2254.7 | 24910 | 3957.1 |
| 1.48E-04 | 54943 | 12775 | 19464 | 3386.9 | 28757 | 4521.9 | 15769 | 2341.8 | 24876 | 3930.5 |
| 2.17E-04 | 55345 | 12792 | 19471 | 3347.7 | 28829 | 4488 | 15879 | 2371.3 | 25107 | 3857.7 |
| 3.18E-04 | 55705 | 12792 | 19457 | 3197.6 | 28904 | 4460.8 | 15911 | 2331.3 | 25186 | 3779.1 |
| 4.67E-04 | 55952 | 12794 | 19597 | 3313.1 | 28969 | 4437.5 | 16028 | 2361.2 | 25254 | 3770 |
| 6.85E-04 | 56123 | 12839 | 19624 | 3177.4 | 29014 | 4445.9 | 16106 | 2340.6 | 25303 | 3753.2 |
| 0.00101 | 56096 | 13003 | 19648 | 3119.1 | 29043 | 4409.6 | 16128 | 2333.5 | 25343 | 3741.5 |
| 0.00148 | 55736 | 13401 | 19651 | 3096.4 | 29042 | 4405.4 | 16112 | 2334.9 | 25322 | 3742.1 |
| 0.00217 | 54678 | 14265 | 19559 | 3105.5 | 28988 | 4420.1 | 16038 | 2338.4 | 25298 | 3746.5 |
| 0.00318 | 52116 | 16268 | 19353 | 3111 | 28796 | 4451.1 | 15903 | 2362.8 | 25155 | 3763.1 |
| 0.00467 | 45955 | 19968 | 18968 | 3176.4 | 28457 | 4494.2 | 15659 | 2401.3 | 24929 | 3786.6 |
| 0.00685 | 35404 | 23068 | 18298 | 3319.6 | 27790 | 4630.5 | 15272 | 2462.9 | 24488 | 3847.3 |

| | | | | | | | | | | |
|---|---|---|---|---|---|---|---|---|---|---|
| 0.0101 | 23595 | 22796 | 17039 | 3692.4 | 26318 | 5053.5 | 14703 | 2578.5 | 23667 | 3985.2 |
| 0.0148 | 14492 | 19468 | 14979 | 4349.4 | 23670 | 5941 | 13797 | 2739.5 | 22264 | 4283 |
| 0.0217 | 8747.9 | 15269 | 11738 | 5326.4 | 19826 | 7009.1 | 12367 | 3063.5 | 19794 | 4992 |
| 0.0318 | 5325.8 | 11472 | 8320.9 | 5563 | 14965 | 7992.7 | 10154 | 3603 | 16048 | 5921.4 |
| 0.0466 | 3289 | 8476.1 | 5582.3 | 5054 | 10618 | 7715.7 | 7544.1 | 3835 | 11877 | 6351.8 |
| 0.0685 | 2054.9 | 6218.5 | 3624 | 4202.2 | 7173.6 | 6581 | 5136.5 | 3651.5 | 8212.3 | 5970 |
| 0.101 | 1312.5 | 4559.4 | 2362.9 | 3326.2 | 4649.4 | 5228.6 | 3376.4 | 3145.7 | 5626.1 | 5116.9 |
| 0.148 | 853.38 | 3347.8 | 1528.4 | 2597.3 | 3069.9 | 4051.8 | 2175.7 | 2496.9 | 3749.9 | 4160.7 |
| 0.217 | 561.09 | 2459.8 | 971.9 | 1945.8 | 1989.2 | 3039.9 | 1378.3 | 1895.8 | 2481.4 | 3250 |
| 0.318 | 364.38 | 1810.8 | 623.13 | 1460.1 | 1256.3 | 2255.1 | 850.79 | 1413.6 | 1604.8 | 2459.3 |
| 0.467 | 238.47 | 1341.2 | 387.11 | 1075.2 | 830.4 | 1642 | 547.34 | 1054 | 983.2 | 1758.8 |
| 0.685 | 150.62 | 991.1 | 229.56 | 778.77 | 546.88 | 1198.2 | 322.06 | 746.71 | 568.5 | 1221.8 |
| 1.01 | 90.698 | 738.74 | 128.98 | 550.59 | 340.61 | 840.83 | 177.12 | 507.88 | 319.06 | 816.48 |
| Amplitude sweep test | Loaded | | | | | | | | | |

| Shear Strain | G' | G" | G' | G" | G' | G" | G' | G" | G' | G" |
|---|---|---|---|---|---|---|---|---|---|---|
| 1 | Pa | Pa | Pa | Pa | Pa | Pa | Pa | Pa | Pa | Pa |
| | T0 | T0 | T1 | T1 | T3 | T3 | T5 | T5 | T7 | T7 |
| 1.00E-04 | 63603 | 14027 | 18375 | 3308 | 16111 | 2652.2 | 23372 | 3862.5 | 15967 | 2553.5 |
| 1.48E-04 | 63894 | 14211 | 18378 | 3293.2 | 16130 | 2628.4 | 23404 | 3884.3 | 16117 | 2554.9 |
| 2.17E-04 | 64451 | 14250 | 18364 | 3268.8 | 16163 | 2652.6 | 23436 | 3826.1 | 16167 | 2496 |
| 3.18E-04 | 64819 | 14143 | 18383 | 3230.6 | 16173 | 2615.4 | 23439 | 3801.6 | 16196 | 2479.6 |
| 4.67E-04 | 65176 | 14081 | 18360 | 3181.3 | 16187 | 2598.5 | 23483 | 3777.1 | 16227 | 2455.1 |
| 6.85E-04 | 65420 | 14100 | 18364 | 3147.7 | 16226 | 2585.4 | 23532 | 3751.5 | 16259 | 2445.7 |
| 0.00101 | 65508 | 14086 | 18361 | 3124 | 16248 | 2579.7 | 23556 | 3758.8 | 16275 | 2437.5 |
| 0.00148 | 65345 | 14213 | 18322 | 3118.2 | 16237 | 2583.1 | 23560 | 3737.9 | 16257 | 2432.7 |
| 0.00217 | 64826 | 14481 | 18245 | 3103 | 16201 | 2582.4 | 23523 | 3745.4 | 16225 | 2429.6 |
| 0.00318 | 63628 | 15131 | 18084 | 3113.1 | 16144 | 2588.8 | 23435 | 3747.1 | 16168 | 2431.5 |
| 0.00467 | 60782 | 16525 | 17834 | 3143.5 | 16016 | 2610.9 | 23259 | 3765.4 | 16053 | 2447.3 |
| 0.00685 | 54347 | 19141 | 17401 | 3206.5 | 15806 | 2642.9 | 22965 | 3783.9 | 15870 | 2472.8 |
| 0.0101 | 42683 | 22089 | 16682 | 3339.1 | 15459 | 2708.8 | 22482 | 3819.3 | 15574 | 2519.7 |

| | | | | | | | | | | |
|---|---|---|---|---|---|---|---|---|---|---|
| 0.0148 | 29427 | 22071 | 15460 | 3628.3 | 14871 | 2824.5 | 21699 | 3898.5 | 15101 | 2587.8 |
| 0.0217 | 18554 | 19178 | 13220 | 4351.3 | 13894 | 3028.6 | 20459 | 4073.7 | 14334 | 2688.1 |
| 0.0318 | 11109 | 15227 | 9845 | 5186.8 | 12367 | 3383.5 | 18444 | 4465.4 | 13090 | 2914.2 |
| 0.0466 | 6618.5 | 11407 | 6542.3 | 5163.3 | 10110 | 3985.9 | 15284 | 5150.2 | 11233 | 3252 |
| 0.0685 | 3946.5 | 8312.6 | 4062.5 | 4468.9 | 7321.7 | 4423.3 | 11411 | 5667.5 | 8938.1 | 3532.1 |
| 0.101 | 2382 | 5970.5 | 2492.9 | 3549.5 | 4792.4 | 4237.7 | 7802.5 | 5472.6 | 6548.3 | 3648.4 |
| 0.148 | 1403.9 | 4214.2 | 1522.5 | 2717.6 | 3005.9 | 3586.7 | 5112.7 | 4712.2 | 4529.3 | 3388.5 |
| 0.217 | 848.68 | 2988.7 | 940.88 | 2036.9 | 1864.4 | 2840.1 | 3302.7 | 3776.1 | 3083.6 | 2849.7 |
| 0.318 | 517.66 | 2125.9 | 593.75 | 1497.7 | 1160.9 | 2177.4 | 2133.5 | 2908.3 | 2060 | 2277.1 |
| 0.467 | 302.68 | 1502.6 | 380.09 | 1110.1 | 723.21 | 1633.1 | 1375.3 | 2189.4 | 1363.2 | 1758 |
| 0.685 | 171.92 | 1055.7 | 239.1 | 812.75 | 458.29 | 1213.8 | 893.77 | 1640.4 | 896.32 | 1322.8 |
| 1.01 | 99.664 | 749.16 | 156.48 | 588.97 | 289.46 | 871.17 | 575.27 | 1208.6 | 595.55 | 977.11 |

**Table S8. Amplitude sweep test on blank and loaded hydrogels immersed in seawater for 7 days.**

| Frequency | Blank |
|---|---|

| sweep test | | | | | | | | | | |
|---|---|---|---|---|---|---|---|---|---|---|
| Angular Frequency | G' | G" | G' | G" | G' | G" | G' | G" | G' | G" |
| rad/s | Pa | Pa | Pa | Pa | Pa | Pa | Pa | Pa | Pa | Pa |
| | T0 | T0 | T1 | T1 | T3 | T3 | T5 | T5 | T7 | T7 |
| 100 | 62621 | 12981 | 25560 | 4138.1 | 46811 | 8071.6 | 29382 | 4530.2 | 23128 | 3264.2 |
| 75 | 60517 | 12662 | 24721 | 4023 | 45353 | 7811 | 28335 | 4360.8 | 22575 | 3162 |
| 56.2 | 58484 | 12379 | 23930 | 3909.4 | 44015 | 7581.7 | 27464 | 4260.5 | 21985 | 3017 |
| 42.2 | 56502 | 12176 | 23211 | 3805.9 | 42752 | 7373.2 | 26691 | 4169.3 | 21455 | 2920 |
| 31.6 | 54613 | 11950 | 22540 | 3701.6 | 41533 | 7186.2 | 25951 | 4069.5 | 20967 | 2844 |
| 23.7 | 52750 | 11850 | 21904 | 3649.5 | 40391 | 7020.2 | 25233 | 4002 | 20508 | 2769.7 |
| 17.8 | 50989 | 11662 | 21280 | 3536.7 | 39302 | 6897.4 | 24550 | 3914.6 | 20069 | 2700.4 |
| 13.3 | 49274 | 11537 | 20674 | 3516.5 | 38227 | 6707.2 | 23935 | 3891.3 | 19658 | 2632.8 |
| 10 | 47571 | 11622 | 20147 | 3410 | 37226 | 6636 | 23284 | 3814.8 | 19245 | 2572 |
| 7.5 | 45897 | 11598 | 19622 | 3403.1 | 36100 | 6582.4 | 22660 | 3826.4 | 18871 | 2525.2 |
| 5.62 | 44284 | 11676 | 19076 | 3377.4 | 35288 | 6390 | 22038 | 3845.2 | 18516 | 2491.7 |
| 4.22 | 42454 | 11889 | 18594 | 3351.3 | 34389 | 6292.7 | 21547 | 3744.8 | 18110 | 2426.3 |

| | | | | | | | | | | |
|---|---|---|---|---|---|---|---|---|---|---|
| 3.16 | 41036 | 11924 | 18188 | 3394.8 | 33526 | 6223.1 | 20991 | 3850.6 | 17796 | 2419.3 |
| 2.37 | 39335 | 12172 | 17476 | 3260.6 | 32676 | 6237.6 | 20575 | 3824 | 17501 | 2354.2 |
| 1.78 | 37819 | 12399 | 17225 | 3322.8 | 31875 | 6190.7 | 20122 | 3918.3 | 17136 | 2462 |
| 1.33 | 36021 | 12549 | 16763 | 3339.4 | 31295 | 5941.9 | 19715 | 3847.1 | 16917 | 2333.6 |
| 1 | 34792 | 12801 | 16582 | 3671.8 | 30617 | 6015.5 | 18929 | 3642.1 | 16528 | 2260.6 |
| 0.75 | 33413 | 12979 | 16240 | 3303.6 | 30036 | 6102 | 18763 | 3491.3 | 16414 | 2335.6 |
| 0.562 | 31956 | 13055 | 16039 | 3435.6 | 29487 | 6081.6 | 18223 | 3397.2 | 16177 | 2264.9 |
| 0.422 | 30778 | 13065 | 15583 | 3220.7 | 29065 | 5698.2 | 18441 | 3696.7 | 15920 | 2279.9 |
| 0.316 | 29150 | 13202 | 15352 | 3035 | 28260 | 6109.1 | 17597 | 3374.5 | 15711 | 2275.6 |
| 0.237 | 27812 | 13212 | 15132 | 3717.6 | 27733 | 5964.9 | 17655 | 3399 | 15420 | 2207.9 |
| 0.178 | 26137 | 13142 | 14496 | 3441.2 | 26829 | 6145.9 | 17625 | 3498.9 | 15260 | 2247.9 |
| 0.133 | 24625 | 13169 | 13943 | 3565.1 | 24724 | 6874.1 | 16201 | 3321.1 | 14835 | 2375.1 |
| 0.1 | 22881 | 13651 | 13628 | 3837.1 | 24346 | 6454.5 | 15782 | 3901.7 | 14770 | 2507.9 |
| Frequency sweep test | Loaded | | | | | | | | | |
| Angular Frequency | G' | G" | G' | G" | G' | G" | G' | G" | G' | G" |

| rad/s | Pa | Pa | Pa | Pa | Pa | Pa | Pa | Pa | Pa | Pa |
|---|---|---|---|---|---|---|---|---|---|---|
| | T0 | T0 | T1 | T1 | T3 | T3 | T5 | T5 | T7 | T7 |
| 100 | 28674 | 7027 | 20043 | 3375.45 | 19402 | 2945.35 | 15377.7 | 2417.6 | 19642 | 3021.7 |
| 75 | 28479 | 6438 | 19326.5 | 3261.55 | 18740.5 | 2806.1 | 14837.5 | 2316.435 | 18973 | 2906.9 |
| 56.2 | 27362.5 | 6882.5 | 18679.5 | 3144.3 | 18161 | 2707.3 | 14337.6 | 2221.21 | 18383 | 2799.8 |
| 42.2 | 27363.5 | 6359 | 18097.5 | 3050.25 | 17628 | 2627.4 | 13917.25 | 2138.125 | 17848.5 | 2728.3 |
| 31.6 | 26498.5 | 6871 | 17546 | 2971.5 | 17131 | 2554.65 | 13522.8 | 2077.32 | 17359 | 2638.8 |
| 23.7 | 26722 | 6450.5 | 17031.5 | 2898.05 | 16656 | 2499.85 | 13158.95 | 2028.15 | 16900.5 | 2570.5 |
| 17.8 | 26102 | 6087 | 16537 | 2830.35 | 16214.5 | 2431.2 | 12836.45 | 1961.07 | 16469.5 | 2518.2 |
| 13.3 | 25532.5 | 5710.5 | 16064 | 2803.55 | 15785 | 2388.75 | 12519.05 | 1936.605 | 16052 | 2457.55 |
| 10 | 25107 | 5362.5 | 15624 | 2768.4 | 15416 | 2349 | 12217.75 | 1883.225 | 15634 | 2418.2 |
| 7.5 | 25736 | 5091.5 | 15191.5 | 2740.55 | 14962 | 2297.15 | 11926.15 | 1856.86 | 15285.5 | 2377.25 |
| 5.62 | 24445 | 4838 | 14785.5 | 2699.3 | 14566.5 | 2253.5 | 11654.55 | 1828.915 | 14918.5 | 2406.7 |
| 4.22 | 24296.5 | 4663.5 | 14392 | 2690.95 | 14118.5 | 2304.8 | 11391.55 | 1789.92 | 14578 | 2323.95 |
| 3.16 | 23131 | 4402 | 13942 | 2780.65 | 13903 | 2230.5 | 11138.15 | 1784.44 | 14214.5 | 2346.25 |
| 2.37 | 24208.5 | 4450 | 13672.5 | 2752 | 13520.5 | 2210.05 | 10898.75 | 1782.39 | 13900 | 2254.5 |

| | | | | | | | | | | |
|---|---|---|---|---|---|---|---|---|---|---|
| 1.78 | 24510 | 4414 | 13298.5 | 2859.7 | 13144.5 | 2230.1 | 10689.7 | 1781.93 | 13585 | 2282.2 |
| 1.33 | 23939 | 4447 | 12936 | 2888.85 | 12813 | 2223.2 | 10470.25 | 1783.835 | 13333 | 2283.7 |
| 1 | 23602 | 4121 | 12584.25 | 2908.95 | 12501 | 2150.1 | 10295.65 | 1791.015 | 13103 | 2224.7 |
| 0.75 | 23261.5 | 4315 | 12356.65 | 2945.6 | 12296 | 2066.2 | 10048.65 | 1761.22 | 12828 | 2300 |
| 0.562 | 23027.5 | 4965 | 12056.3 | 3179.35 | 12069 | 2076.75 | 9738.55 | 1779.63 | 12646 | 2220.95 |
| 0.422 | 22201 | 4463.5 | 11707.05 | 2879.15 | 11824.85 | 2078.1 | 9625.65 | 1615.555 | 12359.5 | 2186 |
| 0.316 | 21817.5 | 4798.5 | 11505.35 | 3095.35 | 11620 | 2026.4 | 9344.85 | 1662.655 | 12169 | 2106.65 |
| 0.237 | 21603 | 4947 | 11272 | 3017.2 | 11315.05 | 1985.55 | 9210.55 | 1830.635 | 11990 | 2111.1 |
| 0.178 | 20981 | 5621 | 10946.4 | 3264.3 | 11113.6 | 2136.8 | 9087.35 | 1755.36 | 11743.5 | 2085.35 |
| 0.133 | 20192 | 5691 | 10062.85 | 3268.15 | 10694.95 | 2086.75 | 8583.55 | 1950.7 | 11457.5 | 2124.55 |
| 0.1 | 19776 | 5532.5 | 9797.05 | 3384.45 | 10320 | 2192 | 7656.55 | 1949.875 | 11108 | 2298.5 |

**Table S9. Frequency sweep test on blank and loaded hydrogels immersed in seawater for 7 days.**

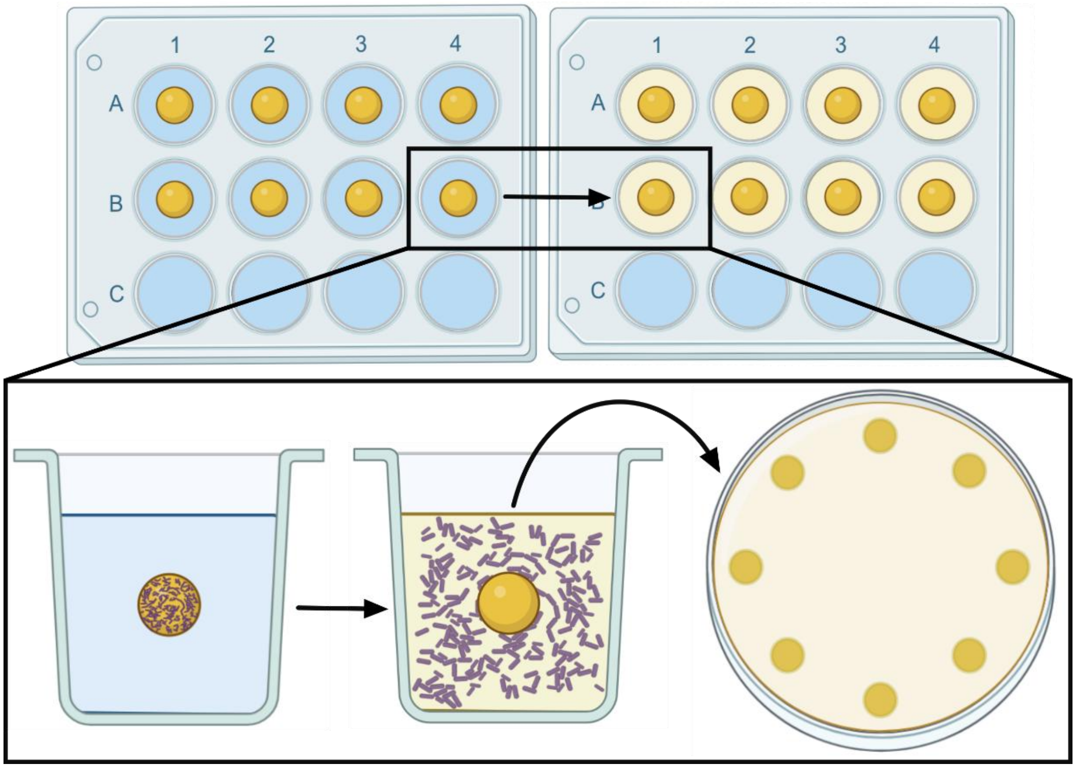

**Figure S5. Probiotic test release *in vitro*.** Schematic illustration of experimental design for probiotic test release *in vitro*.

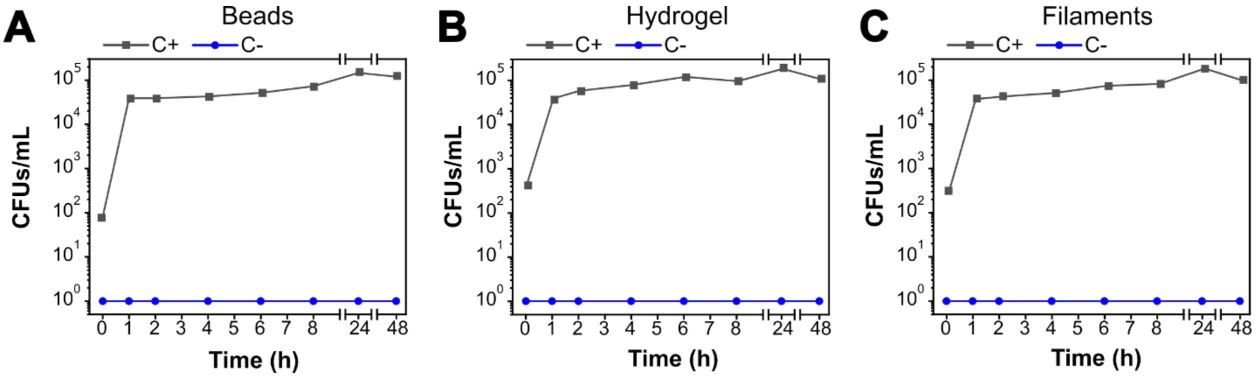

**Figure S6. Release profile of positive and negative controls for the *in vitro* probiotic test release.** (A, B, C) Release curves of bacterial solutions and blank beads, hydrogels, and filaments, respectively.

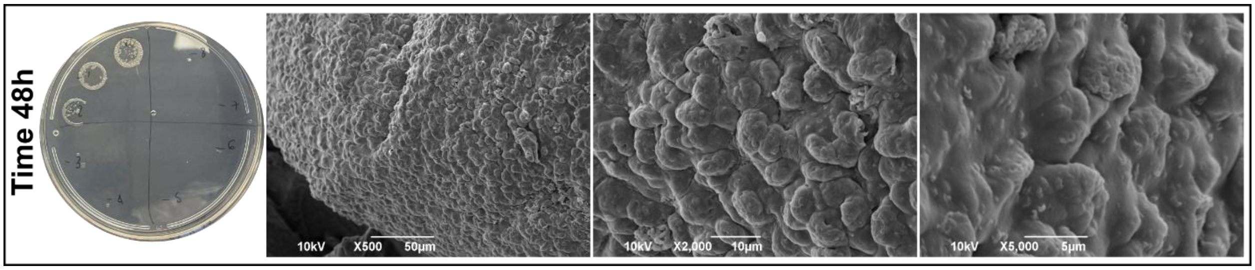

**Figure S7. Probiotic test release *in vitro* at 48h.** Representative photographs of microbial agar plates and SEM images at 48h showing probiotic growth and the surface morphology of the beads.

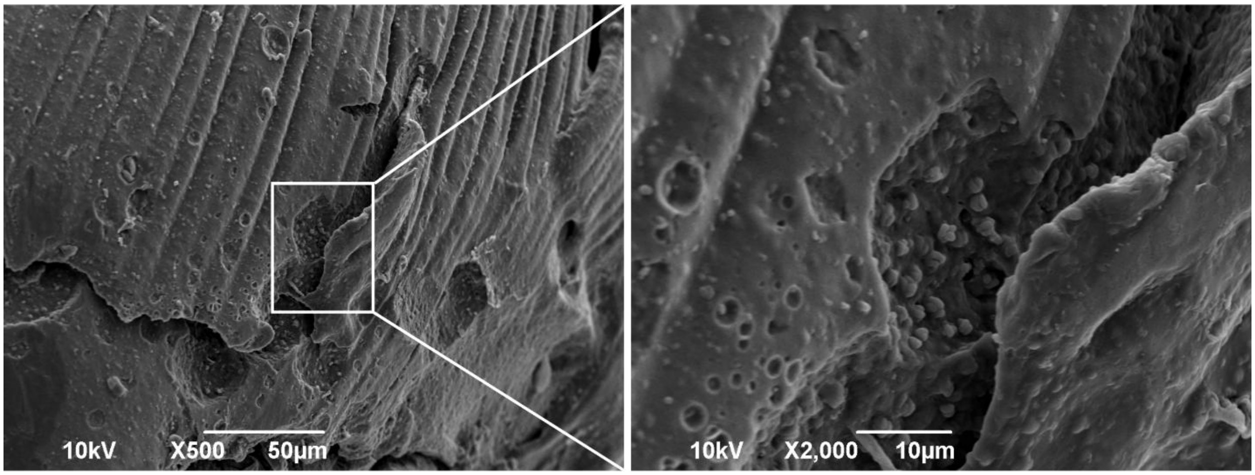

**Figure S8. Cross-section SEM images.** Cross-section SEM image of a loaded bead showing probiotics still entrapped inside the alginate matrix.

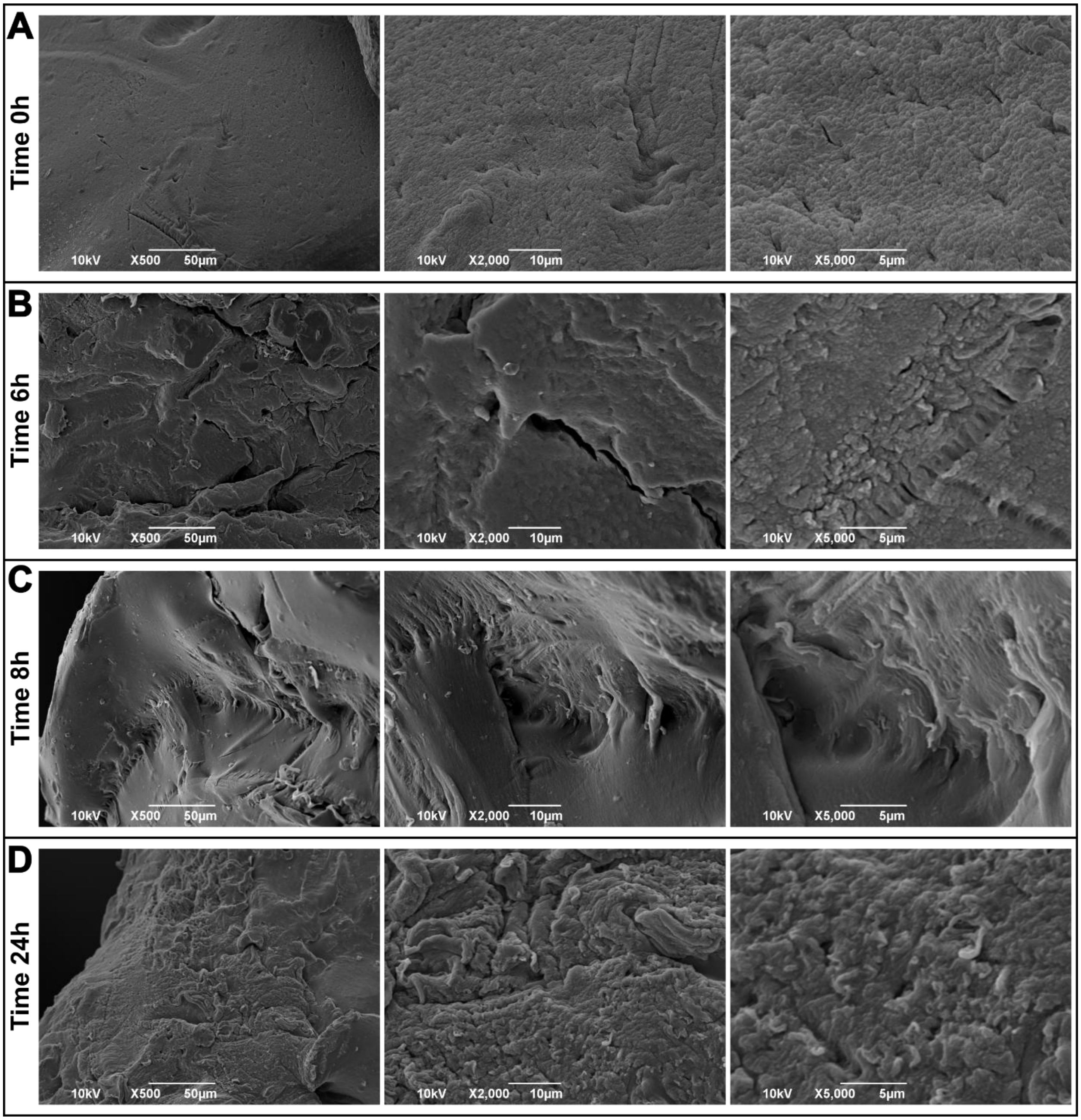

**Figure S9. Control experiment for *in vitro* release.** (A, B, C, and D) Representative SEM images at different time points (0, 6, 8, and 24 hours) showing the surface morphologies of blank beads during the release experiment.

| **Sample** | ***Pseudoalteromonas* relative abundance (%)** |
|---|---|
| **BB-0** | 0 |
| **BB-72** | 27.4 ± 25.2 |
| **BL-0** | 99.9 ± 0.0 |
| **BL-20 min** | 89.8 ± 4.9 |
| **BL-72** | 53.4 ± 11.2 |
| **CT-0** | 2.0 ± 2.2 |
| **CT-72** | 1.0 ± 1.2 |
| **B-0** | 0.0 ± 0.0 |
| **B-8** | 1.4 ± 2.6 |
| **B-24** | 6.2 ± 3.0 |
| **B-48** | 11.5 ± 10.7 |
| **B-72** | 2.1 ± 1.7 |
| **H-0** | 0.3 ± 0.5 |
| **H-8** | 9.3 ± 14.8 |
| **H-24** | 8.3 ± 7.2 |
| **H-48** | 12.7 ± 5.9 |
| **H-72** | 1.7 ± 2.4 |
| **M-0** | 1.5 ± 2.2 |
| **M-8** | 0.3 ± 0.4 |
| **M-24** | 12.6 ± 12.5 |
| **M-48** | 5.0 ± 7.4 |
| **M-72** | 9.4 ± 9.0 |
| **S-0** | 2.4 ± 2.8 |
| **S-8** | 0.8 ± 0.8 |
| **S-24** | 1.6 ± 1.1 |

| | |
|---|---|
| **S-48** | 0.7 ± 0.7 |
| **S-72** | 1.5 ± 2.0 |

**Table S10. Relative abundance of the genus *Pseudoalteromonas* in beads and coral samples.** BB = blank beads; BL = beads loaded with the bacterial strain; CT = control coral; B = coral treated with beads; H = coral treated with hydrogel; M = coral treated with filaments; S = coral treated with syringe injection. Numbers refer to sampling hours. Relative abundances are expressed as average ± st.dev. (n = 4 for bead samples; n = 5 for coral samples).

| BEADS MATERIAL | *PSEUDOALTEROMONAS* | *VIBRIO* | CELLVIBRIONACEAE |
|---|---|---|---|
| **BB-0.1** | Not detected | Not detected | Not detected |
| **BB-0.2** | Not detected | Not detected | Not detected |
| **BB-0.3** | Not detected | Not detected | Not detected |
| **BB-0.4** | Not detected | Not detected | Not detected |
| **BB-72.1** | 3.49156532 | 27.31463319 | 38.27971754 |
| **BB-72.2** | 11.71297222 | 30.24185489 | 33.02018789 |
| **BB-72.3** | 34.8184655 | 14.34444499 | 39.52895141 |
| **BB-72.4** | 59.41564745 | 1.543389633 | 35.3038245 |
| **BL-0.1** | 0 | 0 | 0 |
| **BL-0.2** | 0 | 0 | 0 |
| **BL-0.3** | 0 | 0 | 0 |
| **BL-0.4** | 0 | 0 | 0 |
| **BL-20 MIN.1** | 99.97252575 | 0 | 0 |
| **BL-20 MIN.2** | 99.9771907 | 0 | 0 |
| **BL-20 MIN.3** | 99.96212394 | 0 | 0 |
| **BL-20 MIN.4** | 99.88409032 | 0 | 0 |
| **IIT-BL-72.1** | 69.21591348 | 13.90498262 | 12.25376593 |
| **IIT-BL-72.2** | 51.07649568 | 16.65839865 | 23.53408076 |
| **IIT-BL-72.3** | 42.8150442 | 18.67164477 | 30.87632985 |
| **IIT-BL-72.4** | 50.3803132 | 28.49117574 | 12.89087745 |

| CORAL CTRL | *PSEUDOALTEROMONAS* | *VIBRIO* | CELLVIBRIONACEAE |
|---|---|---|---|
| **CT-0.1** | 1.614205004 | 0 | 0.484261501 |
| **CT-0.2** | 0.889903177 | 1.559018541 | 0.164071678 |
| **CT-0.3** | 5.658496752 | 1.38940593 | 0.08406086 |
| **CT-0.4** | 1.673306773 | 0 | 0 |
| **CT-0.5** | 0.078595756 | 0.890751899 | 0 |

| | | | |
|---|---|---|---|
| **CT-72.1** | 0.217904485 | 0 | 0.744506991 |
| **CT-72.2** | 0.940986692 | 0.645248017 | 1.236725366 |
| **CT-72.3** | 0.922431866 | 0.109014675 | 0.067085954 |
| **CT-72.4** | 0 | 5.682715075 | 0 |
| **CT-72.5** | 2.988215488 | 0.252525253 | 0.126262626 |

| CORAL BEADS | *PSEUDOALTEROMONAS* | *VIBRIO* | CELLVIBRIONACEAE |
|---|---|---|---|
| **B-0.1** | 0 | 0 | 0 |
| **B-0.2** | 0.050889146 | 2.849792203 | 0 |
| **B-0.3** | 0 | 0 | 0 |
| **B-0.4** | 0 | 0 | 0 |
| **B-0.5** | 0 | 0 | 0 |
| **B-8.1** | 0.571428571 | 0 | 0 |
| **B-8.2** | 6.064281383 | 10.61249242 | 0 |
| **B-8.3** | 0 | 0 | 0 |
| **B-8.4** | 0 | 0.766417001 | 0 |
| **B-8.5** | 0.314349112 | 5.824704142 | 0 |
| **B-24.1** | 1.56424581 | 0.893854749 | 0 |
| **B-24.2** | 4.944308612 | 2.091822874 | 0 |
| **B-24.3** | 6.969059778 | 3.574647041 | 0 |
| **B-24.4** | 9.041720063 | 4.172006265 | 0 |
| **B-24.5** | 8.238871993 | 4.147083218 | 0 |
| **B-48.1** | 30.16591252 | 6.033182504 | 0 |
| **B-48.2** | 3.529071449 | 0.202487706 | 6.754411339 |
| **B-48.3** | 6.269966514 | 1.835957046 | 8.41768985 |
| **B-48.4** | 9.770114943 | 6.781609195 | 2.298850575 |
| **B-48.5** | 7.616487455 | 2.956989247 | 0.201612903 |
| **B-72.1** | 3.783068783 | 5.873015873 | 0 |
| **B-72.2** | 2.221371122 | 6.600280863 | 0 |
| **B-72.3** | 0.588019111 | 6.321205439 | 0 |
| **B-72.4** | 0.113004991 | 4.030511348 | 0 |
| **B-72.5** | 3.585007275 | 5.146012684 | 0 |

| CORAL HYDROGEL | *PSEUDOALTEROMONAS* | *VIBRIO* | CELLVIBRIONACEAE |
|---|---|---|---|
| **G-0.1** | 0.540008027 | 3.043018207 | 0 |
| **G-0.2** | 1.071428571 | 0 | 0 |
| **G-0.3** | 0.075832701 | 3.424138132 | 0 |
| **G-0.4** | 0 | 0 | 0 |
| **G-0.5** | 0 | 0 | 0.767754319 |
| **G-8.1** | 2.871818819 | 3.829091758 | 0 |
| **G-8.2** | 1.40625 | 1.40625 | 0 |
| **G-8.3** | 3.93442623 | 3.114754098 | 0 |

| | | | |
|---|---|---|---|
| **G-8.4** | 2.637362637 | 7.692307692 | 0 |
| **G-8.5** | 35.81700967 | 0.607488222 | 0 |
| **G-24.1** | 4.705030764 | 58.34238147 | 1.121968874 |
| **G-24.2** | 4.010938924 | 15.06381039 | 0.47857794 |
| **G-24.3** | 20.52055407 | 51.63650581 | 0.228999106 |
| **G-24.4** | 3.166437215 | 10.43764945 | 0.432939642 |
| **G-24.5** | 9.118773946 | 30.34482759 | 0.536398467 |
| **G-48.1** | 17.2528363 | 3.517017828 | 7.852512156 |
| **G-48.2** | 19.97105644 | 7.525325615 | 0 |
| **G-48.3** | 11.82943604 | 3.370013755 | 0 |
| **G-48.4** | 8.833413346 | 14.97839654 | 0.38406145 |
| **G-48.5** | 5.69604714 | 1.211228415 | 0.229151322 |
| **G-72.1** | 0.844482881 | 4.024011802 | 0.330671008 |
| **G-72.2** | 0.4247053 | 5.705770925 | 0.036196474 |
| **G-72.3** | 6.02006689 | 1.170568562 | 0.836120401 |
| **G-72.4** | 0.568053676 | 1.030390167 | 0 |
| **G-72.5** | 0.694783383 | 6.342051621 | 0.172260343 |

**CORAL FILAMENTS**

| | ***PSEUDOALTEROMONAS*** | ***VIBRIO*** | **CELLVIBRIONACEAE** |
|---|---|---|---|
| **M-0.1** | 0 | 1.005025126 | 0 |
| **M-0.2** | 5.34591195 | 0.314465409 | 0 |
| **M-0.3** | 0.296880323 | 1.348128681 | 0.158173943 |
| **M-0.4** | 0.503778338 | 0 | 0 |
| **M-0.5** | 0.620017715 | 0 | 0 |
| **M-8.1** | 0.787888151 | 3.958751738 | 0 |
| **M-8.2** | 0 | 3.993392927 | 0 |
| **M-8.3** | 0 | 0 | 0 |
| **M-8.4** | 0 | 0 | 0 |
| **M-8.5** | 0.72426612 | 3.454685297 | 0 |
| **M-24.1** | 9.991435912 | 27.43362832 | 1.769911504 |
| **M-24.2** | 1.694527135 | 8.003205862 | 0.274788184 |
| **M-24.3** | 27.50664011 | 34.9435591 | 3.569057105 |
| **M-24.4** | 23.56828194 | 17.73127753 | 3.414096916 |
| **M-24.5** | 0.257069409 | 9.254498715 | 0 |
| **M-48.1** | 18.13471503 | 4.317789292 | 0 |
| **M-48.2** | 0.102272222 | 0.100048913 | 0.015563164 |
| **M-48.3** | 2.161263508 | 6.733167082 | 2.992518703 |
| **M-48.4** | 2.182600676 | 3.842606824 | 2.951122041 |
| **M-48.5** | 2.329009434 | 4.76120283 | 1.533018868 |
| **M-72.1** | 8.463949843 | 37.72204807 | 5.538140021 |
| **M-72.2** | 3.932151118 | 47.87972244 | 3.315343099 |
| **M-72.3** | 3.243670886 | 27.76898734 | 4.272151899 |
| **M-72.4** | 25.04708098 | 6.779661017 | 0 |

| | | | |
|---|---|---|---|
| **M-72.5** | 6.220095694 | 27.13602187 | 8.339029392 |

| CORAL SYRINGE | *PSEUDOALTEROMONAS* | *VIBRIO* | CELLVIBRIONACEAE |
|---|---|---|---|
| **S-0.1** | 0.141094379 | 0.443805957 | Not detected |
| **S-0.2** | 1.706910908 | 1.186511241 | Not detected |
| **S-0.3** | 1.58778626 | 5.06870229 | Not detected |
| **S-0.4** | 0.896336711 | 3.039750585 | Not detected |
| **S-0.5** | 0.395517469 | 3.724456163 | Not detected |
| **S-8.1** | 0 | 0.618556701 | Not detected |
| **S-8.2** | 0.248756219 | 0 | Not detected |
| **S-8.3** | 0.565504241 | 0.329877474 | Not detected |
| **S-8.4** | 1.944065484 | 0.409276944 | Not detected |
| **S-8.5** | 1.040781648 | 2.113423959 | Not detected |
| **S-24.1** | 1.869530628 | 3.102625298 | Not detected |
| **S-24.2** | 1.602513747 | 2.892772977 | Not detected |
| **S-24.3** | 1.234567901 | 0 | Not detected |
| **S-24.4** | 0 | 0.686813187 | Not detected |
| **S-24.5** | 3.175630615 | 2.342983798 | Not detected |
| **S-48.1** | 1.121794872 | 2.083333333 | Not detected |
| **S-48.2** | 0 | 3.954132068 | Not detected |
| **S-48.3** | 1.56626506 | 2.530120482 | Not detected |
| **S-48.4** | 0.570484912 | 1.621378171 | Not detected |
| **S-48.5** | 0 | 0 | Not detected |
| **S-72.1** | 0 | 0 | Not detected |
| **S-72.2** | 0.112641725 | 0.585367655 | Not detected |
| **S-72.3** | 4.892354209 | 1.101654582 | Not detected |
| **S-72.4** | 0.940438871 | 0 | Not detected |
| **S-72.5** | 1.426533524 | 0 | Not detected |

**Table S11. Relative abundance of key bacterial taxa.** Relative abundance (%) of *Pseudoalteromonas* sp., *Vibrio* sp., and members of the family Cellvibrionaceae within the total microbiome of each sample analyzed.

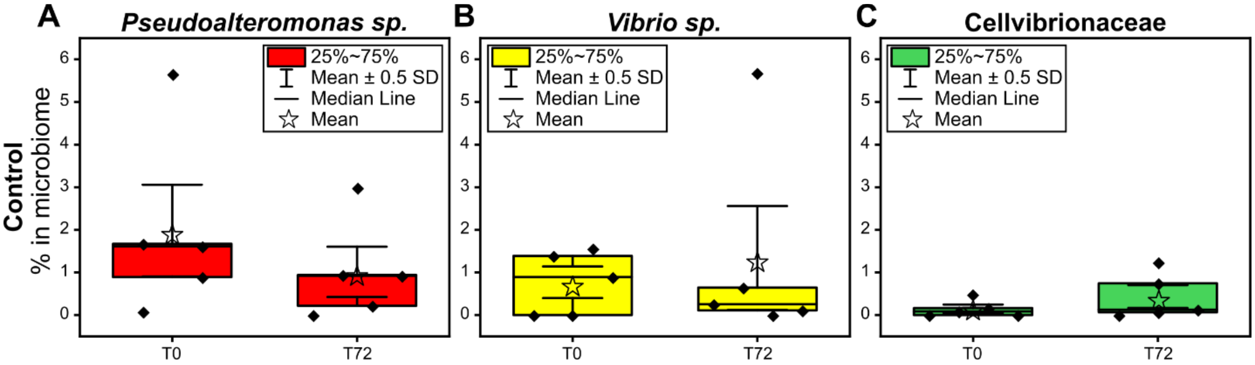

**Figure S10. Relative abundance (%) of *Pseudoalteromonas* spp., *Vibrio* spp., and unclassified Cellvibrionaceae within the coral microbiome of control corals.** (A, B, and C) Relative abundance of *Pseudoalteromonas* spp., *Vibrio* spp., and unclassified Cellvibrionaceae on control corals at the beginning and at the end of the in vivo experiment, respectively.

| Group 1 | Group 2 | p-adjusted |
|---|---|---|
| **H-48** | B-0 | 0.055 |
| **H-48** | H-0 | 0.056 |
| **H-48** | B-8 | 0.056 |
| **H-48** | M-8 | 0.056 |
| **B-24** | B-0 | 0.056 |
| **H-48** | M-0 | 0.056 |
| **H-48** | S-8 | 0.056 |
| **S-48** | H-48 | 0.056 |
| **CT-72** | H-48 | 0.056 |
| **S-72** | H-48 | 0.056 |
| **H-72** | H-48 | 0.056 |
| **B-24** | H-0 | 0.059 |
| **B-24** | M-8 | 0.059 |
| **H-48** | CT-0 | 0.061 |

| | | |
|---|---|---|
| **H-48** | S-24 | 0.061 |
| **H-48** | S-0 | 0.063 |
| **S-48** | B-24 | 0.063 |
| **B-72** | H-48 | 0.063 |
| **B-24** | S-8 | 0.064 |
| **CT-72** | B-24 | 0.068 |

**Table S12. Two-sample t-test** pairwise comparisons that were considered only marginally not significant (adjusted p-values ranging 0.055-0.068). CT = control coral; B = coral treated with beads; H = coral treated with hydrogel; M = coral treated with filaments; S = coral treated with syringe injection. Numbers refer to sampling hours.

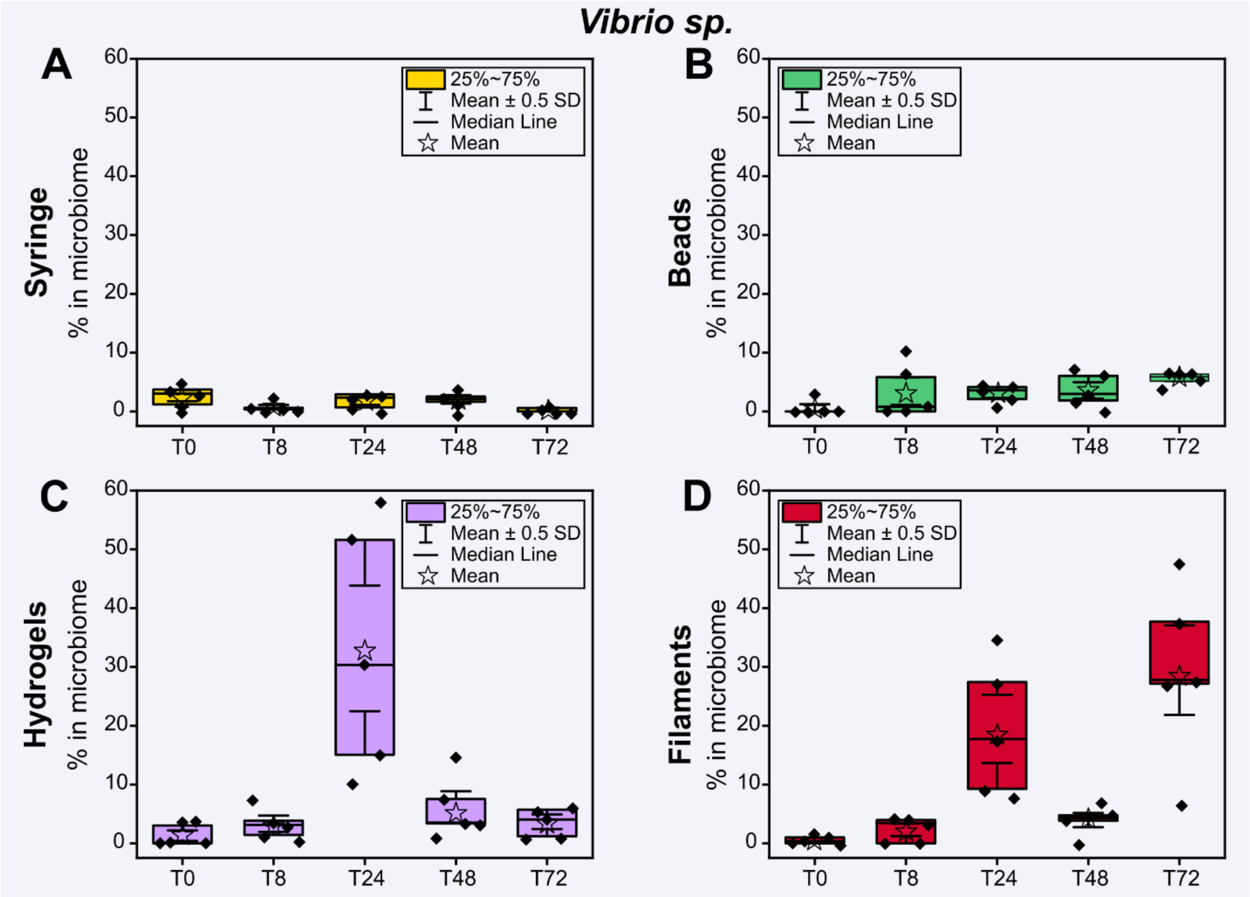

**Figure S11. Relative abundance (%) of *Vibrio* spp. within the coral microbiome during the four in vivo treatments.** (A, B, C, and D) Relative abundance of *Vibrio* spp. during the syringe, beads, hydrogel, and filament treatments, respectively.

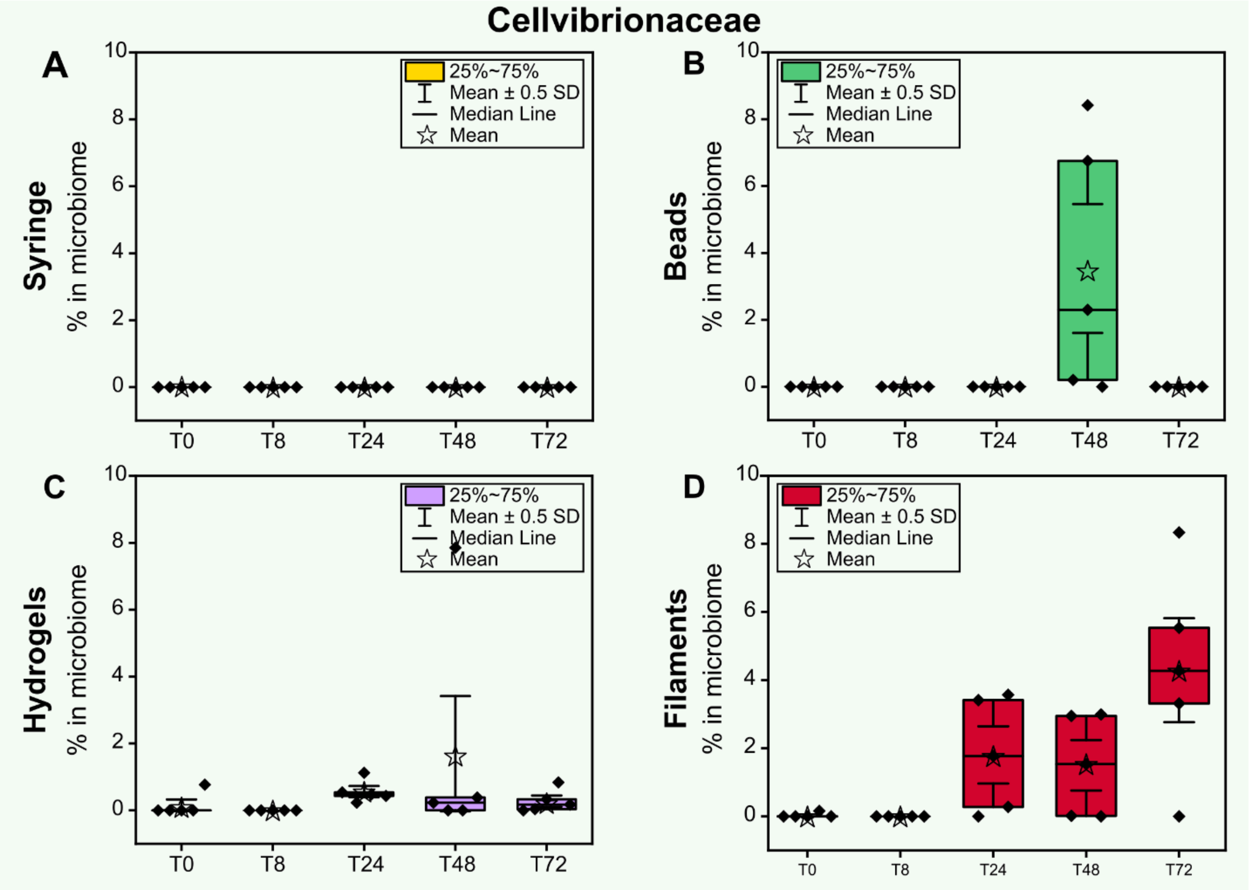

**Figure S12. Relative abundance (%) of species belonging to the family Cellvibrionaceae within the coral microbiome during the four in vivo treatments.** (A, B, C, and D) Relative abundance of members of the family Cellvibrionaceae during the syringe, beads, hydrogel, and filament treatments.

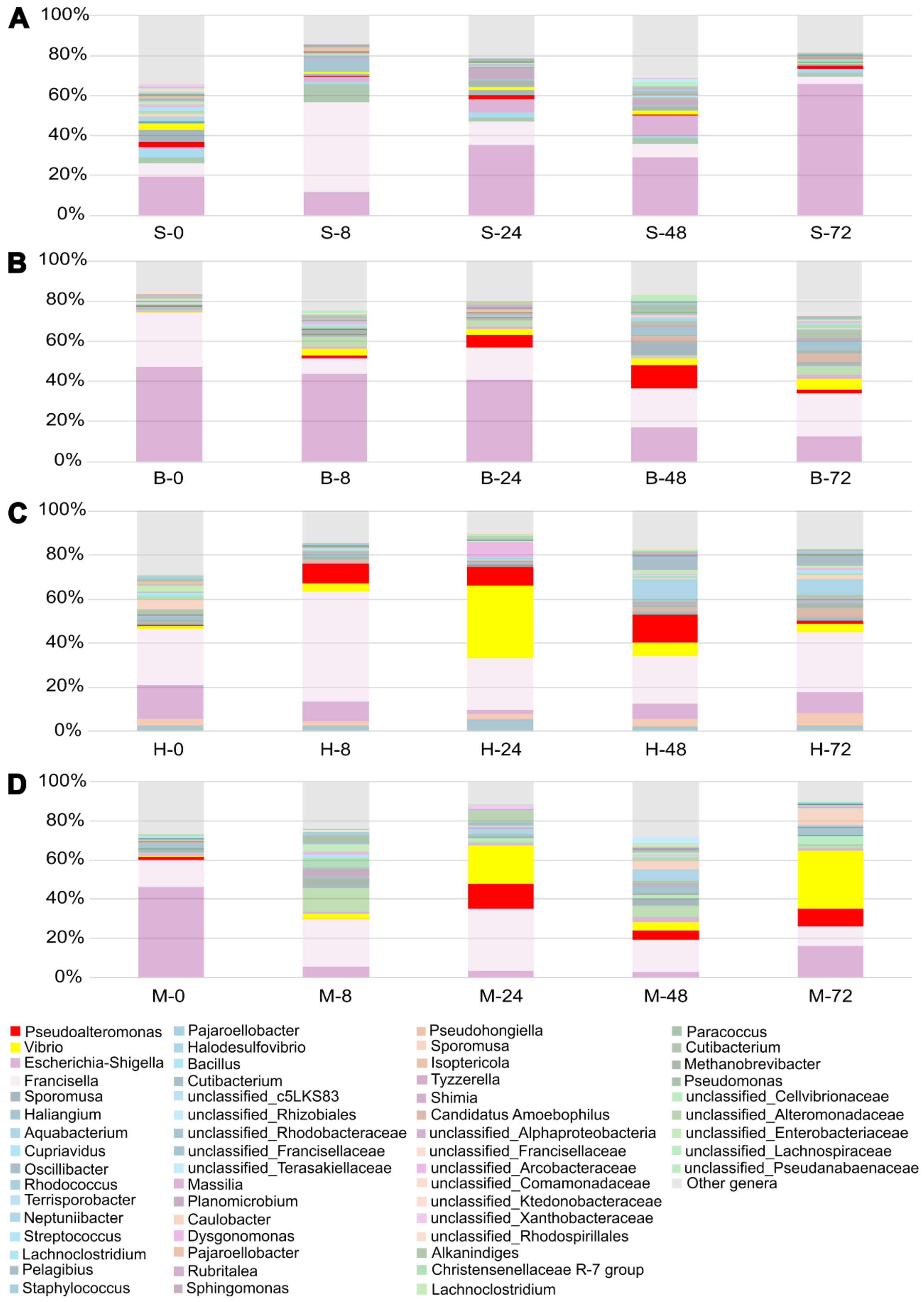

**Figure S13. Microbiome analysis**. (A) syringe, (B) beads, (C) hydrogels, and (D) filaments.

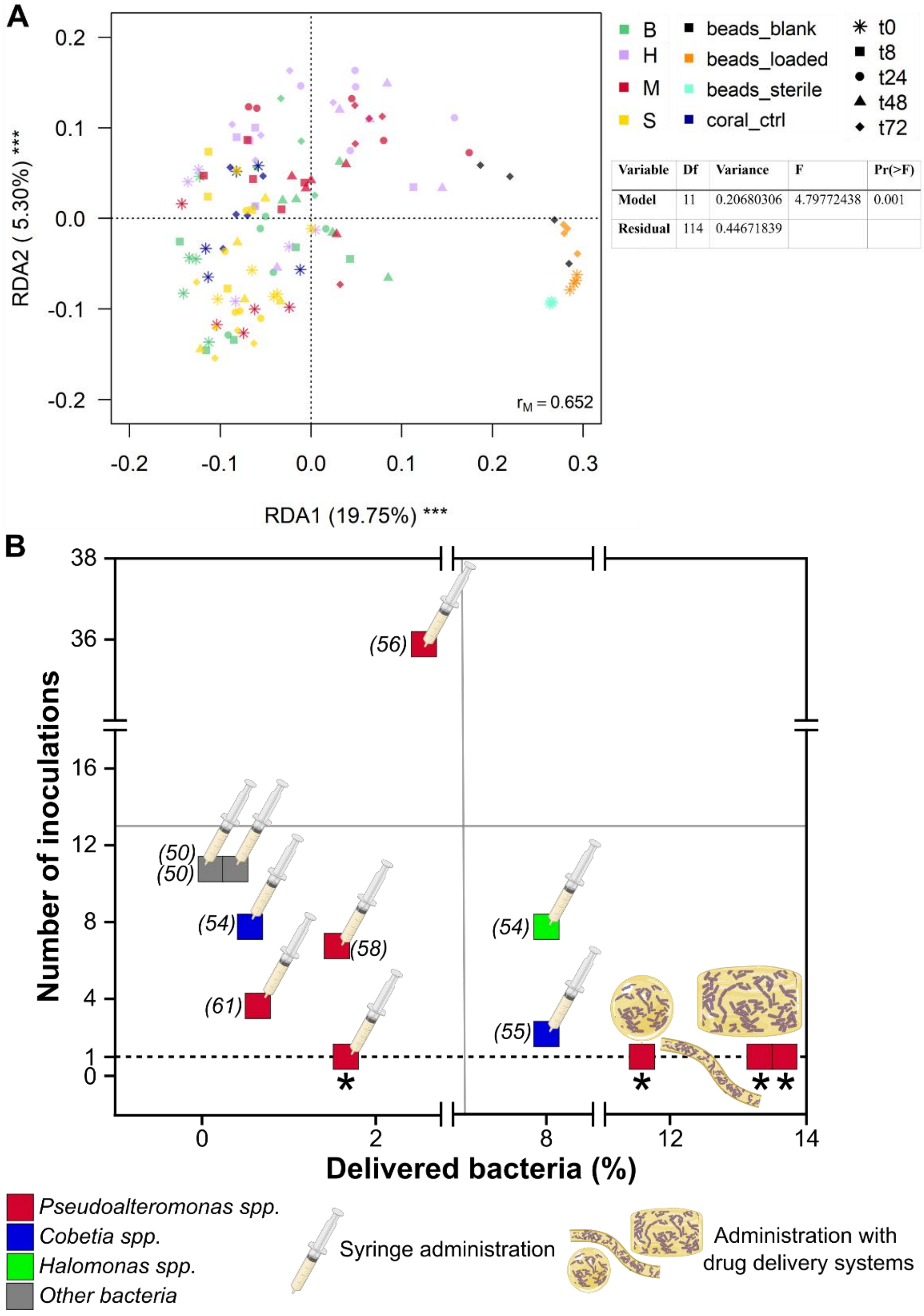

| Variable | Df | Variance | F | Pr(>F) |
|---|---|---|---|---|
| Model | 11 | 0.20680306 | 4.79772438 | 0.001 |
| Residual | 114 | 0.44671839 | | |

**Figure S14. RDA and schematic representation of data available in literature.** (A) Redundancy analysis (RDA) of coral-associated microbial communities during the *in vivo* experiment under different alginate-based treatments. RDA1 and RDA2 explained 19.75% and 5.30% of the total variance, respectively. Samples are colored by treatment group, and symbols indicate sampling time points (0, 8, 24, 48, and 72 hours). (B) Schematic summary of reported probiotics inoculation and integration into the coral microbiome, highlighting the number and via (syringe, beads, hydrogels, and filaments) of inoculations, and the bacterial concentrations used or achieved across published studies. Inoculation of *Pseudoalteromonas spp.* is shown as red squares, C*obetia spp.* as blue squares, *Halomonas spp.* as green, and other bacteria as grey. Red squares marked with an asterisk refer to the results presented in this work.

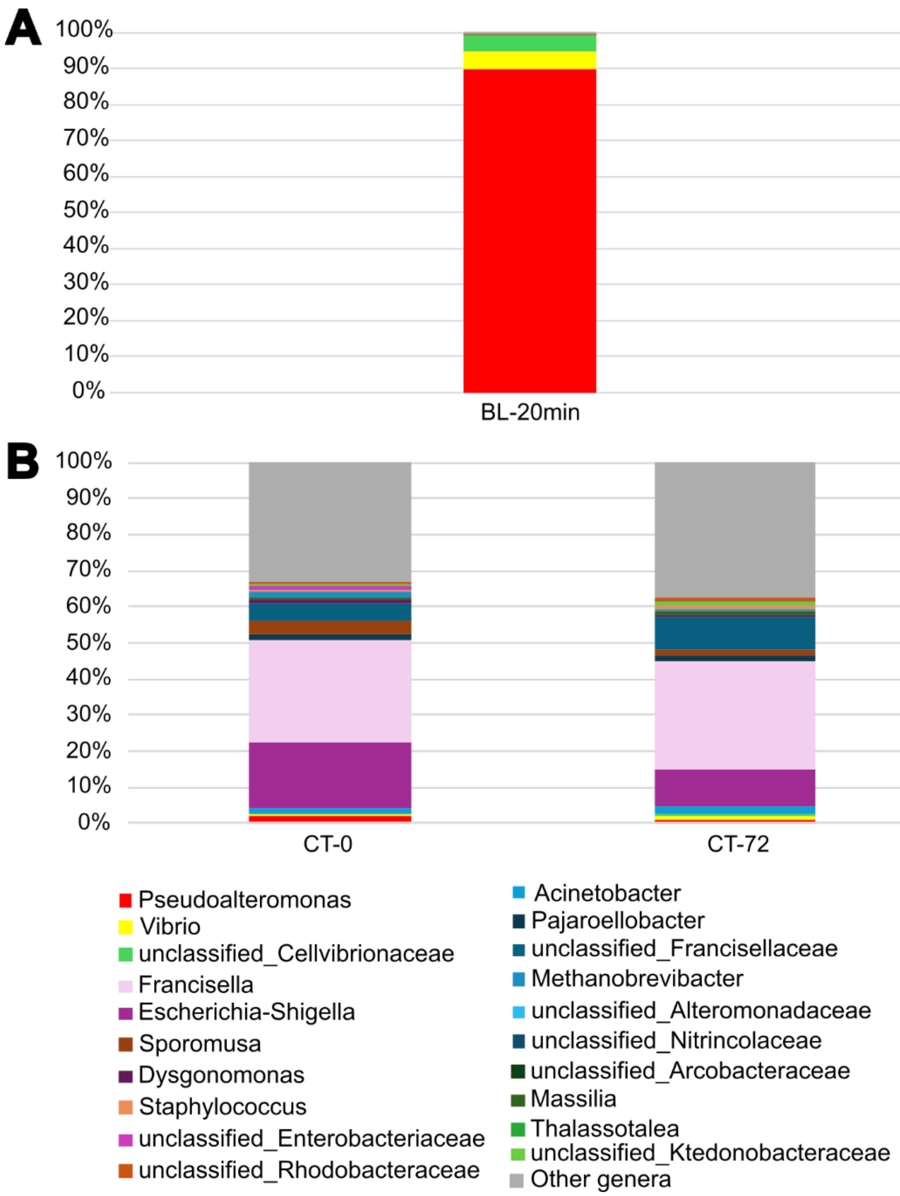

**Figure S15. Microbiome analysis on control corals and loaded beads.** (A) Microbiome analysis on 4 blank beads sampled 20 minutes after being in contact with the coral, to see when the biological colonisation started. (B) Microbiome analysis on control corals.

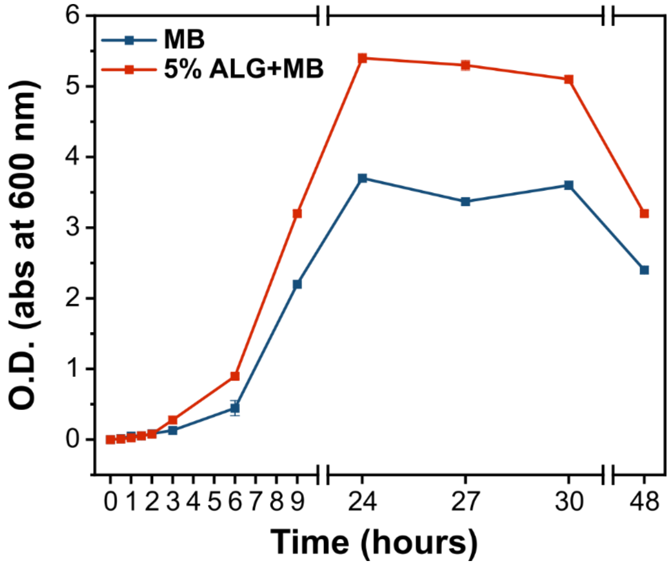

**Figure S16.** ***Pseudoalteromonas peptidolytica*** **growth curve performed in MB and in 5% alginate solution.** The growth curve followed a standard four-stage growth pattern anticipated for bacteria, with a lag phase followed by the exponential phase at the beginning, the stationary phase in which the plateau is reached, and finally the death phase. The highest O.D. at 600nm was observed after 24 hours.

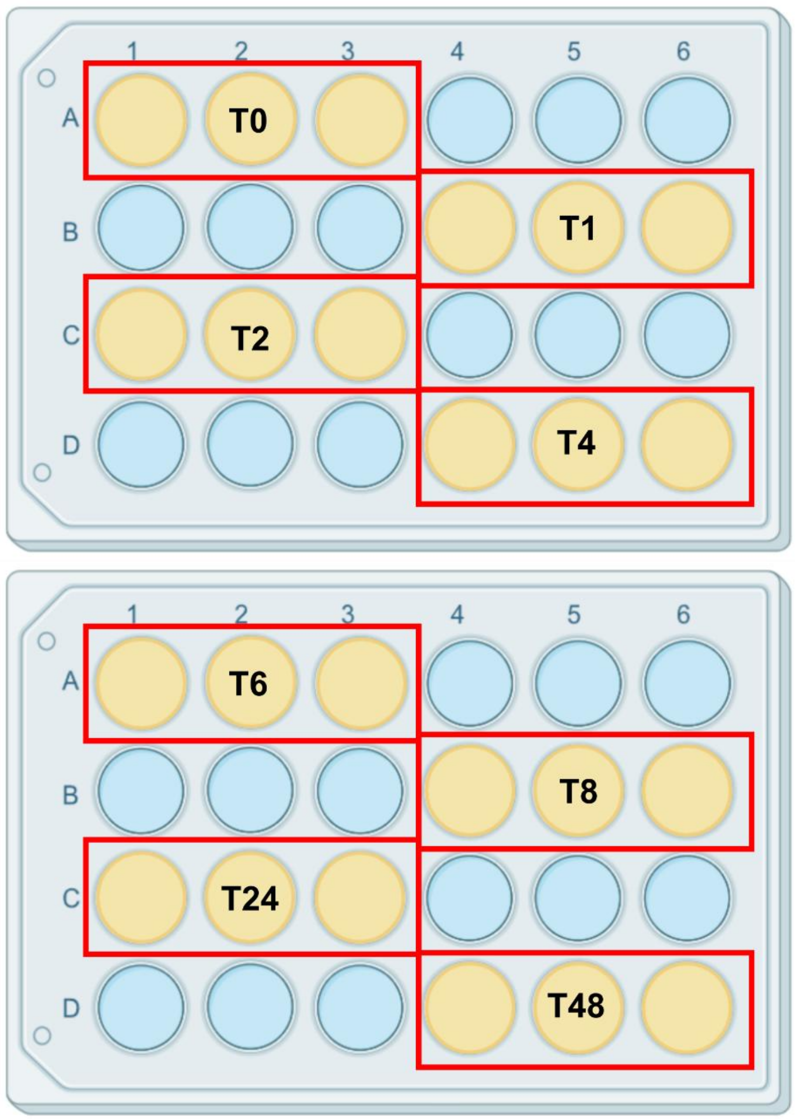

**Figure S17. Schematic representation of the sampling procedure for bacterial release.** Samples were collected from three wells at 0, 1, 2, 4, 6, 8, 24, and 48 h and subjected to serial dilutions ranging from $10^{-1}$ to $10^{-8}$ before being plated in triplicate.

### Illumina sequencing results

Illumina sequencing of prokaryotic communities associated with coral samples was used both to check successful delivery of the probiotic strain and to evaluate the overall composition of coral microbiomes. After quality filtering and removal of non-prokaryotic ASVs, one single ASV (sequence:
CCGCGGTAACACCGGCAGCTCTAGTGGTAGCCATTTTTATTGGGCCTAAAGCGTTCGT
AGCCGGTTTAATAAGTCTCTGGTGAAATCCTGTAGCTTAACTATTGGAATTGCTGGAG
ATACTATTAGACTTGAGGTCGGGAGAGGTTAGAGGTACTCCCAGGGTAGGGGTGAAA
TCCTGTAATCCTGGGAGGACCACCTGTGGCGAAGGCGTCTAACTGGAACGAACCTGA
CGGTGAGGGACGAAAGCTAGGGGCGCGAACCGG) was recognized as an accidental laboratory contamination and was therefore discarded from the dataset. A total of 2,118,097 valid sequences were obtained, ranging between 531 and 148,106 per sample. All samples were retained in subsequent analyses.
